# Cis-regulatory mutations associate with transcriptional and post-transcriptional deregulation of the gene regulatory program in cancers

**DOI:** 10.1101/2020.06.25.170738

**Authors:** Jaime A. Castro-Mondragon, Miriam Ragle Aure, Ole Christian Lingjærde, Anita Langerød, John W. M. Martens, Anne-Lise Børresen-Dale, Vessela Kristensen, Anthony Mathelier

## Abstract

**Background:** Most cancer alterations occur in the noncoding portion of the human genome, which contains important regulatory regions acting as genetic switches to ensure gene expression occurs at correct times and intensities in correct tissues. However, large scale discovery of noncoding events altering the gene expression regulatory program has been limited to a few examples with high recurrence or high functional impact.

**Results:** We focused on transcription factor binding sites (TFBSs) that show similar mutation loads than what is observed in protein-coding exons. By combining cancer somatic mutations in TFBSs and expression data for protein-coding and miRNA genes, we evaluated the combined effects of transcriptional and post-transcriptional alteration on the dysregulation of the regulatory programs in cancer. The analysis of seven cancer cohorts culminated with the identification of protein-coding and miRNA genes linked to mutations at TFBSs that were associated with a cascading trans-effect deregulation on the cells’ regulatory program. Our analyses of cis-regulatory mutations associated with miRNAs recurrently predicted 17 miRNAs as pan-cancer-associated through deregulation of their target gene networks. Overall, our predictions were enriched for protein-coding and miRNA genes previously annotated as cancer drivers. Functional enrichment analyses highlighted that cis-regulatory mutations are associated with the dysregulation of key pathways associated with carcinogenesis

**Conclusions:** These pan-cancer results suggest that our method predicts cis-regulatory mutations related to the dysregulation of key gene regulatory networks in cancer patients. It highlights how the gene regulatory program is disrupted in cancer cells by combining transcriptional and post-transcriptional regulation of gene expression.

## INTRODUCTION

Dysregulation of the gene expression regulatory program in a cell is a hallmark of cancer. The often observed aberrant gene expression in cancer can be triggered by deregulation at any regulatory level (transcriptional and post-transcriptional) [1,2]. While a majority of studies have focused on somatic mutations lying within protein-coding regions, most alterations occur in the noncoding portion of the human genome, which contains cis-regulatory elements acting as genetic switches to ensure gene expression occurs at correct times and intensities in the correct cells and tissues [3]. Molecular alterations at these regions can alter the regulatory network of the cells, conferring oncogenic behaviours, which has been associated with clinical and histopathological features in cancer [3]. However, identification of noncoding cancer driver events at cis-regulatory regions has been limited to a few examples with high recurrence or high functional impact [3–7]. In recent work based on mutation recurrence along the human genome, the Pan-Cancer Analysis of Whole Genomes (PCAWG) consortium, claimed that patients harbour ~4.6 driver mutations. The PCAWG consortium estimated that driver point mutations in noncoding regions (~1.2 per patient) were less frequent than driver point mutations in protein-coding genes (~2.6 per patient) [8]. However, large scale discovery of noncoding drivers has been hindered by their low level of recurrence, the target size of functional elements, technical shortcomings, and their composite effect with small individual effect size on multiple regulatory regions, e.g. slightly altering, but not obliterating, protein-DNA interactions [4,8]. Further, while high-impact driver mutations are typically sought, medium-impact putative passengers can have an aggregated effect in tumorigenesis, beyond annotated driver events [9].

Gene expression is mainly regulated at the transcriptional level by the binding of transcription factors (TFs) to promoters (cis-regulatory regions surrounding genes’ transcription start sites, TSSs) and enhancers (cis-regulatory regions distal to genes) at TF binding sites (TFBSs) [10,11]. Most of the studies that predict noncoding driver mutations in cis-regulatory regions rely on the identification of mutational hotspots, which are regions with higher mutation frequencies than expected by chance [8,12–18]. Other studies explore somatic mutations with potential effect on TF-DNA interactions [19–22] based on DNA sequence information alone, followed by *in vitro* experiments to confirm the potential impact of the predicted mutations on gene expression. Other studies directly combine somatic mutation data with gene expression information to evaluate the impact of the mutations in cancer samples. For instance, studies identified differential allele-specific expression of genes between cancer and normal cells to pinpoint causal cis-regulatory variations in breast cancers [23,24]. Mutations close to the TSSs of genes were shown to exert in-cis effect on the expression of the corresponding genes [25]. Another example is the *xseq* tool that associates mutations with changes in expression in gene networks [26]. The tool has been originally developed to predict mutations in protein-coding exons with trans-effect [26] and adapted to consider noncoding mutations associated with protein-coding genes in B cell lymphomas [27]. This methodology specifically assesses the trans-associations between mutations and gene network expression alteration in cancer samples through either exonic or cis-regulatory mutations linked to protein-coding genes [26,27].

At the post-transcriptional level, miRNAs control gene expression by acting as ‘buffers’ to induce translational repression and mRNA degradation [28,29]. miRNA biogenesis generally comprises three steps in mammals: transcription of a primary transcript (pri-miRNA) that can be several kilobases long, cleavage of the pri-miRNA into a precursor (pre-miRNA) of ~70bp, and cleavage of the precursor to produce mature miRNAs of ~22bp [29,30]. The mature miRNA sequence is then loaded in the RNA-induced silencing complex to specifically target mRNAs for repression through base pair complementarity at the 3’UTR of mRNA targets. A miRNA sequence is predicted to target tens to hundreds of mRNAs [31]. The influence of miRNA-based regulation on mRNA translation is not an on/off system but rather an interplay between miRNA-binding site specificity and miRNA abundance [28,32]. Therefore, even small changes in miRNA abundance may have an effect on the expression of several direct targets but also other mRNAs through a cascading effect, potentially leading to dysregulation patterns observed in cancer. This observation, amongst others, suggests that miRNAs can act as cancer drivers [33,34].

Despite active research on post-transcriptional regulation and the identification of miRNAs and their targets [35], the understanding of miRNA transcriptional regulation is currently limited [30]. One obstacle was the lack of precise identification of pri-miRNA TSSs. The FANTOM5 consortium recently took advantage of the cap analysis of gene expression (CAGE) technology to identify pri-miRNA TSSs genome-wide from different cell types and tissues in human and mouse [36]. Given their short size and the fact that they are not recurrently mutated [8], we hypothesize that the driver potential of miRNAs in cancer could be driven by cis-regulatory mutations that alter their expression in cancers with downstream cascading effect on the gene regulatory program of the cells.

The recent availability of high-quality sets of direct TF-DNA interactions [37], miRNA TSS locations [36], somatic cancer mutations, and cancer cell expression data [38] provides an unprecedented opportunity to analyze alterations of gene regulatory programs in cancer by looking at both transcription and post-transcriptional levels of gene expression regulation. The PCAWG consortium stated that the community is facing a ‘paucity’ in the discovery of noncoding cancer drivers that could be shortened by analyzing larger sample datasets [8]. We hypothesize that focusing on regulatory variants within TFBSs associated with protein-coding and miRNA genes combined with gene expression data has the potential to pinpoint cis-regulatory variants linked to the dysregulation of key gene regulatory networks in cancer patients.

In this study, we adapted the framework of the *xseq* tool to predict cis-regulatory somatic mutations associated with the dysregulation of gene networks by considering both protein-coding and miRNA genes. We predicted genes that are associated with cis-regulatory mutations and cascading trans-effects on the gene regulatory program alteration across seven cancer patient cohorts from The Cancer Genome Atlas (TCGA) [38]. This analysis revealed 17 miRNAs recurrently predicted in the different cohorts. Functional enrichment analyses of the deregulated networks confirmed that pathways known to be associated with carcinogenesis are recurrently disrupted. We conclude that interpretation of noncoding mutations can be improved by focusing on TF-DNA interactions with the combined analysis of both transcriptional and post-transcriptional regulation of gene expression to revert the paucity in the discovery of cancer-associated noncoding events.

## RESULTS

### Transcription factor binding sites harbour similar mutational load than protein-coding exons

We considered somatic mutations from whole genome sequencing of 349 samples from seven cancer patient cohorts (35 to 92 samples per cohort) covering seven distinct cancer types from TCGA [38] (Additional file 1). Specifically, we selected samples where trios of somatic mutations, RNA-seq, and small RNA-seq data were available. In aggregate, we examined 11,434,931 somatic single nucleotide variants and small insertions and deletions (from 2,832 to 1,014,969 per sample; Additional file 2; Figure S1).

To highlight cancer-associated cis-regulatory mutations, we considered a set of TFBSs predicted as direct TF-DNA interactions in the human genome and stored in the UniBind database [37]. We first assessed whether this set of TFBSs would represent regions of functional interest similar to the coding portion of the human genome commonly studied to predict cancer-associated mutations. These TFBSs cover ~2.2% (68,071,257 nt) of the human genome, close to the exonic coverage of protein-coding genes (~2.6%; 81,416,464 nt). Focusing on the somatic mutations, we observed that 1-2% of the mutations in each sample are lying within these TFBSs (median of 277 mutations per sample; Additional file 2; Figure S2). As expected, mutation rates in TFBSs varied between cancer cohorts but were similar to the mutation rates observed in exons of protein-coding genes (two-tailed Wilcoxon tests p-values between 0.13 and 0.96; Figures 1 and S3-S4). We observed that TFBSs were less mutated than their surrounding sequences with mutation rates increasing as the size of the flanking regions increases (Figure 1). Note that regions of 1 kb surrounding TFBSs harbour mutation rates similar to what is expected by chance (two-tailed Wilcoxon tests p-values between 0.56 and 0.95; Figures S3-S4). While exons exhibit mutation rates similar to those observed within TFBSs (Figure S5), their flanking regions show a smaller increase in mutation rates than the increase detected in the vicinity of TFBSs (Figures S6-S7).

**Figure 1.**
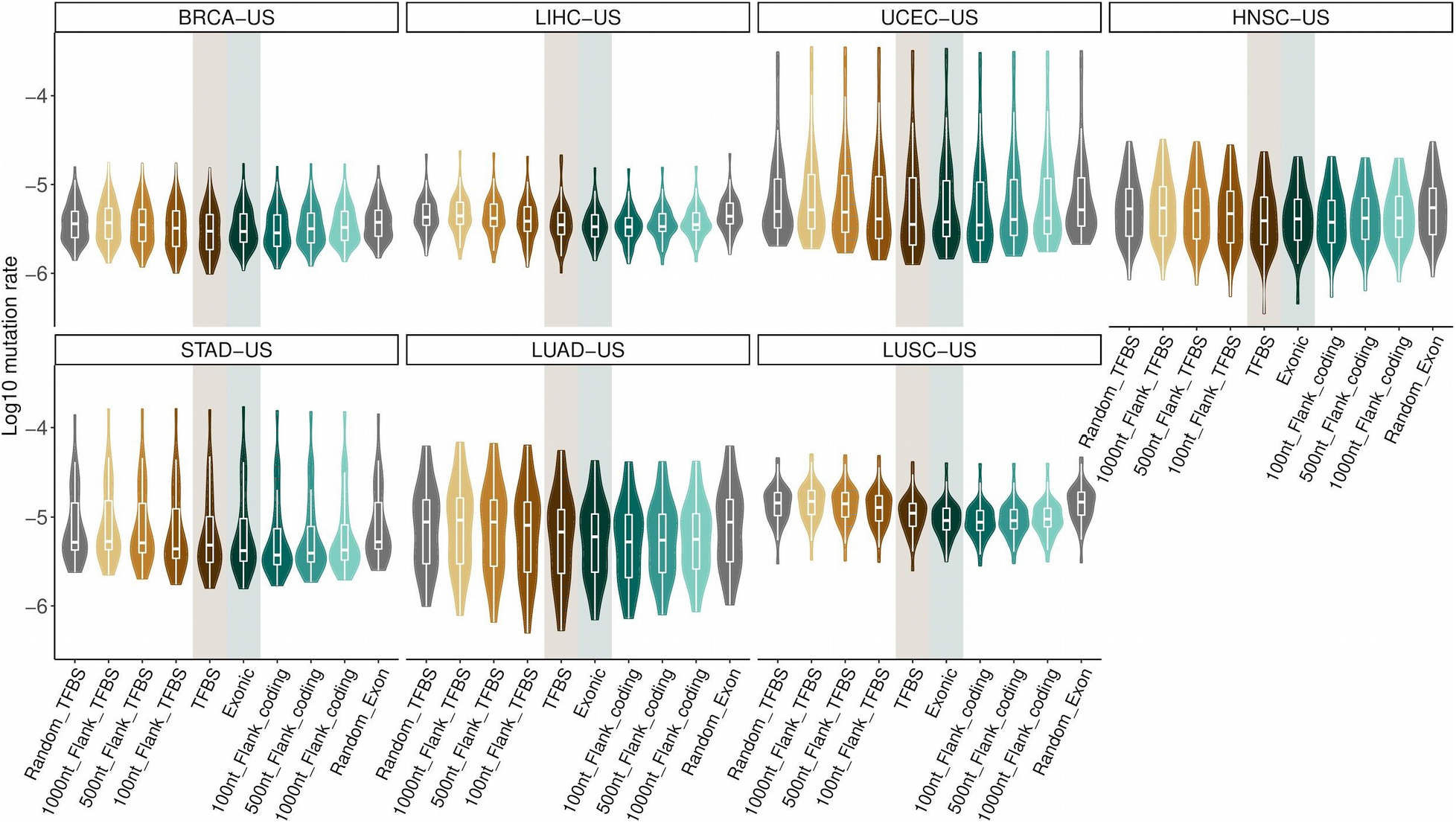
Comparison of mutation rates in TFBSs and exons versus their flanking regions and random mutation rates. Each panel corresponds to a specific cancer cohort (see title boxes) and each point corresponds to a sample. On each panel, the two central boxplots (shadowed) represent mutation rates in TFBS and exonic regions, the remaining box plots correspond to mutation rates in increasing-size flanking regions (100, 500, and 1000 nt) and mutation rates expected by chance (150 randomly distributed sets of mutations in the genome; Material and methods).

Taken together, these results highlight that the mutation frequencies in the studied set of TFBSs follow a similar pattern to what is observed in protein-coding exons. It provides an a posteriori confirmation that the set of TFBSs we considered is likely composed of functional regions in the human genome and could be used to highlight cis-regulatory mutations of functional interest in cancer genomes.

### Cis-regulatory and loss-of-function mutations are complementary mechanisms to alter protein-coding gene networks

We sought to predict the cis-regulatory mutations lying in these TFBSs and that were linked to cascading effects on gene network deregulation, a hallmark of carcinogenic events. We first focused on the mutations linked to protein-coding genes and compared their effect to mutations altering the function of protein-coding genes. Specifically, we considered a protein-coding gene to be mutated through either a loss-of-function (LoF) somatic mutation in one of its exons as in ref. [26] or a somatic mutation overlapping a TFBS associated with the gene. TFBSs were linked to protein-coding or miRNA genes based on cis-regulatory element-to-gene associations from GeneHancer [39] or distances to TSSs (Material and methods; Figure S8). We related the mutations to their potential trans-effect on expression disruption in protein-coding gene networks using the *xseq* tool, following approaches implemented in previous studies [26,27]. Specifically, the method uses a hierarchical bayesian approach to associate mutations with expression dysregulation in biological networks associated with the mutated protein-coding genes. In a nutshell, it assesses the posterior probability of the likely association between observing mutations in a set of patients and observed deviations from neutral expression in these samples for protein coding genes in the same network. The likely trans-associations between mutations and gene network deregulation are first assessed in a sample-specific manner and then across samples from the same cohort (Figure S9). Genes with low expression in a given cohort were filtered out; the distribution of the 90th percentile of expression for genes was decomposed into two Gaussian distributions corresponding to low and high expression values and only genes lying in the high expression distribution were conserved (Material and methods). Further, we corrected for copy number alteration to compensate for their cis-effect on expression (Material and methods). LoF mutations and mutations overlapping TFBSs were analyzed independently.

Pan-cancer analyses of the seven TCGA cohorts predicted 30 protein-coding genes when considering LoF mutations (none in HNSC-US, LUAD-US, and LUSC-US; 2 in LIHC-US; 4 in BRCA-US; 9 in STAD-US; 18 in UCEC-US) and 283 genes when considering cis-regulatory mutations (6 in LIHC-US; 22 in BRCA-US and HNSC-US; 35 in LUSC-US; 42 in STAD-US; 81 in LUAD-US; 107 in UCEC-US) (Figures 2 and S10-S12). Three genes were linked to dysregulated networks in association with both LoF and cis-regulatory mutations but in different patients and cohorts: *ACVR2A*, *ARID1A*, and *GATA3* (Figure 2A). These three genes are already known cancer drivers that we predict to be impacted by alternative mutational mechanisms (LoF or cis-regulatory mutations). For the other genes, we observed that they were either associated with LoF mutations or mutations in TFBSs across cohorts (e.g. *TP53*, *RPL22*, and *PDS5B* with LoF mutations; *PIK3C3* and *CEBPB* with cis-regulatory mutations; Figure 2).

**Figure 2.**
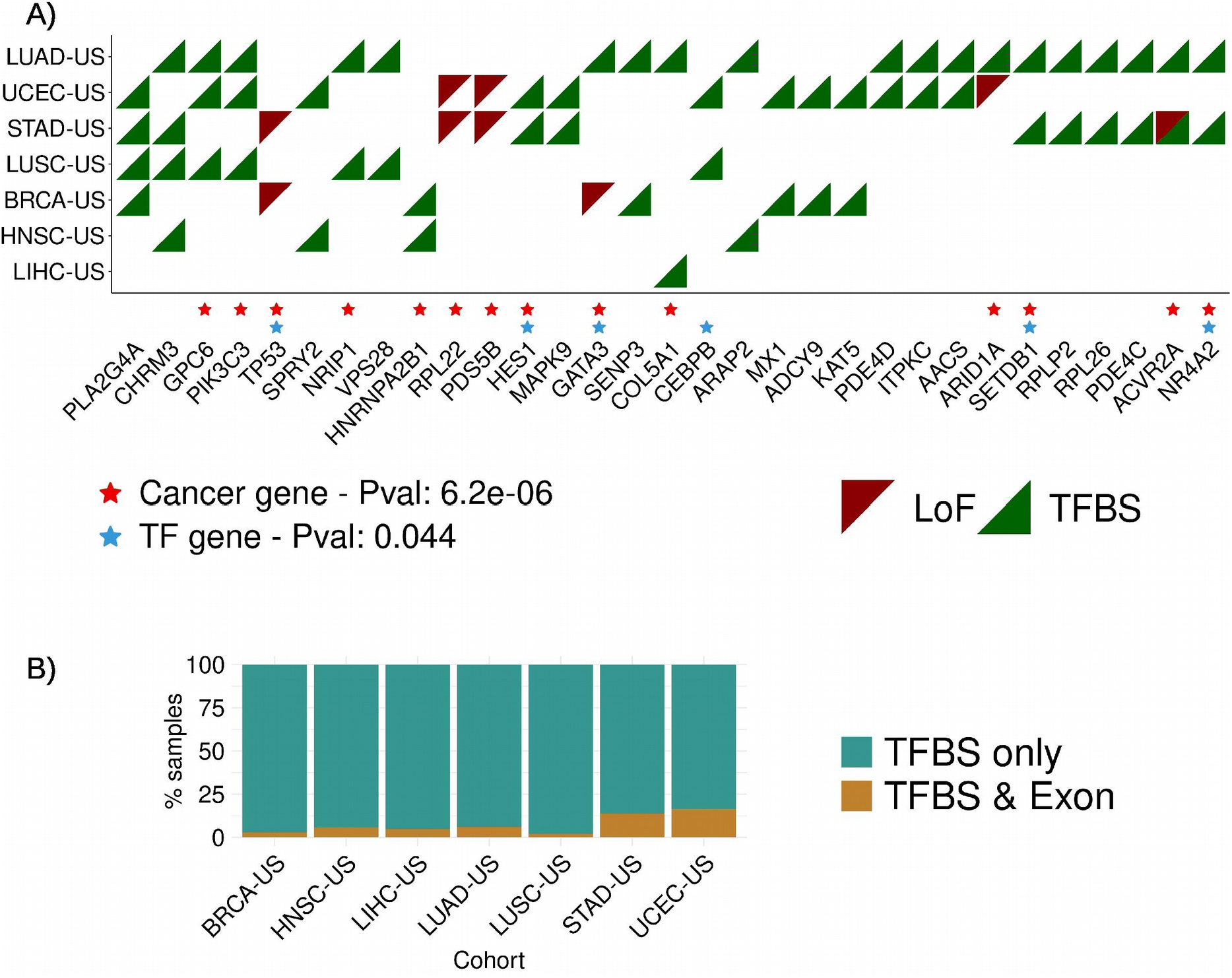
Pan-cancer predicted protein-coding genes. **A)** Predictions were obtained applying the *xseq* tool when considering protein-coding genes mutated through either LoF (red triangles) or cis-regulatory (TFBS; green triangles) mutations, independently. Genes predicted in at least two cohorts are depicted here. Genes known as cancer genes (red stars) and TFs (blue stars) were found to be enriched (hypergeometric tests; p-values provided in the legend; Material and methods). **B)** Samples where genes were predicted through cis-regulatory mutations were considered for each cohort and assessed for the presence of LoF mutations in the same genes for the same cohort (TFBS & Exon) or no LoF mutation in the corresponding gene (TFBS only).

From the combined list of 310 predicted protein-coding genes (Additional Files 3 and 4), 87 were already annotated as cancer genes (p-value = 1.5e-16; hypergeometric test) and 37 as TFs (p-value = 0.0061; Figures S10-S12). Considering recurrent predictions over cancer types, we observed 31 genes to be predicted in at least two cohorts. These 31 genes are enriched for already known cancer drivers (p-value = 6.2e-6; hypergeometric test) and TFs (p-value = 0.044; hypergeometric test) (Figure 2A).

The genes identified through cis-regulatory mutations rarely contained LoF mutation in the same patients (Figures 2B and S13). These results reinforce the possible complementary mechanisms between LoF and cis-regulatory mutations at play in cancer patients to alter the gene regulatory program of cancer cells. We observed that multiple genes could be predicted in the same sample through cis-regulatory mutations (e.g. from 1 to 42 genes were predicted in UCEC-US samples). Nevertheless, these genes tend to be interconnected in the dysregulated genes’ networks with a maximum of 5 disconnected subgraphs per sample (Figure 3). All these genes are predicted through mutations associated with cascading trans-effect in gene network dysregulation but the method cannot pinpoint which specific event could be the main driver event or if it is due to the combination of cis-regulatory mutations. When considering all the predicted genes per cohort, we detect a similar pattern with subnetworks of interconnected genes (Figures 3B and S14). Altogether, these interconnections suggest that the predicted genes are likely involved in similar biological pathways with altered expression associated with cis-regulatory somatic mutations.

**Figure 3.**
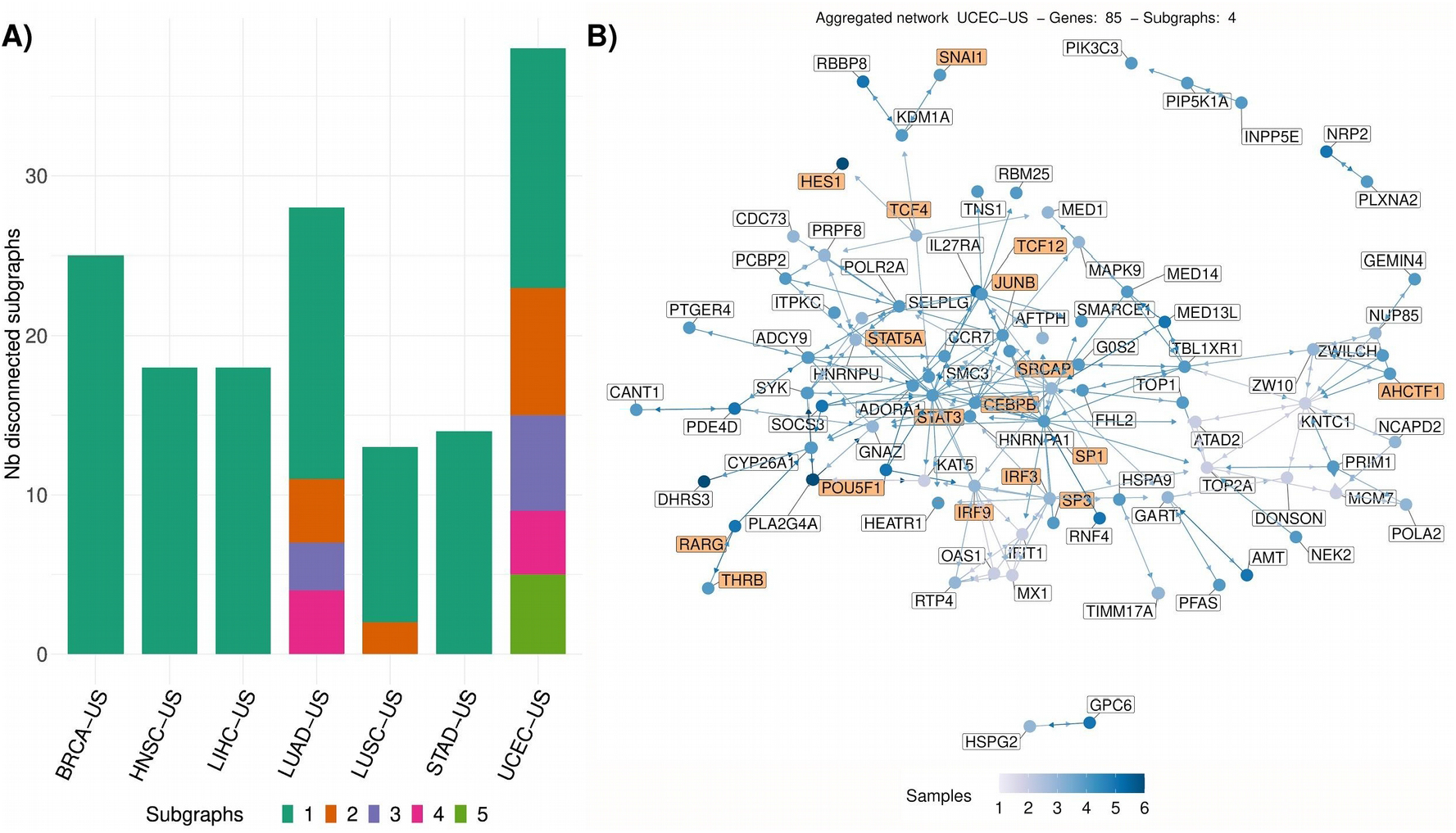
Networks of predicted protein-coding genes. **A)** Stacked histogram depicting the number of disconnected networks of predicted protein-coding genes (see legend) per sample (number of samples on the y-axis) for each cohort (x-axis). **B)** Network of all predicted genes in the UCEC-US cohort. The number of samples in which each gene was predicted is provided using a color scale (see legend). TF genes are highlighted with an orange background.

### Deregulation of transcriptional activity and cancer pathways are trans-effect signature of the predicted cis-regulatory and loss-of-function mutations

Next, we performed enrichment analyses to shed light on the functional role of the somatic mutations predicted to be associated with a cascading effect on gene expression alteration. One advantage of *xseq* is its capacity to highlight the specific genes in the biological networks associated with the candidate cancer-associated genes that are dysregulated in the samples harbouring the somatic mutations considered (Material and methods) [26]. We observed that these genes were consistently found to be either up- or down-regulated in the samples with predicted disrupted expression (see the blue and red colors in the upper and lower clusters in Figure 4A). These results highlight sets of genes up- or down-regulated across samples where cancer-associated genes are predicted.

**Figure 4.**
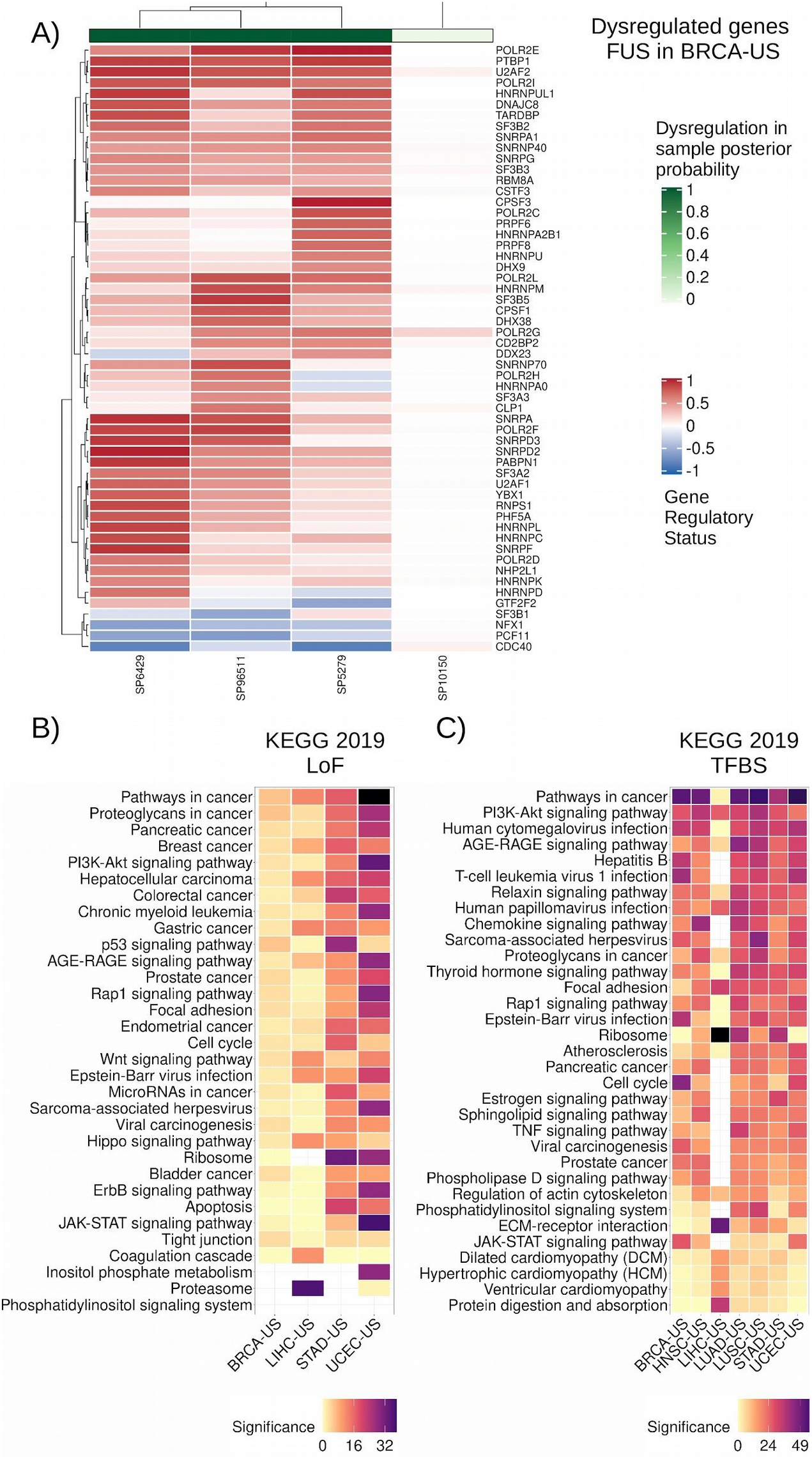
Dysregulated protein-coding gene networks and functional enrichment analysis. **A)** Dysregulated gene network in samples where FUS is predicted through cis-regulatory mutations in BRCA-US (rows: dysregulated genes associated with FUS; columns: samples with FUS-associated cis-regulatory mutations). The color scale represents the gene regulatory status posterior probability (red: up-regulation; blue: down-regulation). The green horizontal bar on top shows the sample-specific dysregulation posterior probability computed by *xseq* for the samples harboring a cis-regulatory mutation in the FUS gene. **B)** KEGG 2019 most enriched terms computed from all the dysregulated genes associated with the predicted protein-coding genes (Figure 4A is one example for GATA3) by *xseq* with LoF and **C)** cis-regulatory mutations in TCGA cohorts (columns). Terms (rows) are ordered by their mean rank across all cohorts. Significance is provided as −log_10_(p-value).

We assessed the biological relevance of the networks predicted to be dysregulated in association with the protein-coding genes predicted through LoF or cis-regulatory mutations. Functional enrichment analysis was performed using pathways from KEGG [40], WikiPathways [41], and Panther [42], and gene ontology biological processes (GO BP [43]) with the EnrichR tool [44]. The dysregulated genes in the networks are enriched for transcriptional activity (‘regulation of transcription, DNA−templated’ from GO BP; Figure S15). Combined with the enrichment of TFs in the list of predicted cancer-associated genes, this result emphasizes that the alteration of transcriptional regulation is a common feature of cancer cells throughout cancer types. Focusing on biological pathways enriched in our list of genes from the dysregulated networks, we found pathways already known to be associated with carcinogenesis at the top of the enriched terms (e.g. ‘Pathways in cancer’, ‘WNT signaling’, ‘PI3K-Akt signaling’, and ‘Focal adhesion’; Figures 4B-C and S15-S18). These results confirm that our approach highlighted somatic exonic and cis-regulatory mutations associated with potential protein-coding cancer-associated genes with cascading effect on regulatory alteration of key cancer-related pathways.

The enrichment for cancer pathways represents a posteriori confirmation that our method can pinpoint somatic events likely associated with carcinogenesis. Nevertheless, our results suggest that alteration of gene network expression in different patients could be achieved through cis-regulatory mutations associated with different genes involved in the same pathway.

### Combining transcriptional and post-transcriptional regulation highlights pan-cancer miRNAs associated with gene expression alteration

The analysis of protein-coding genes presented above exhibited that our methodology is able to pinpoint cis-regulatory mutations likely associated with carcinogenesis. With miRNAs involved in post-transcriptional regulation of gene expression, we hypothesized that our method could highlight cis-regulatory mutations linked to miRNAs with downstream cascading effect on the gene regulatory program of the cells. This new analysis aimed at combining transcriptional (through mutations in TFBSs) and post-transcriptional (through miRNA-targets regulatory networks) regulation to predict miRNAs associated with a trans-effect on gene expression alteration through somatic mutations in cis-regulatory elements.

Pan-cancer analyses of the seven TCGA cohorts predicted 98 miRNAs derived from 63 pre-miRNAs as potential cancer-associated miRNAs (Figure S19; Additional Files 3 and 4). From these 98 miRNAs, 73 were already annotated as cancer miRNAs in the miRCancer database [48] (p-value = 9.6e-28; hypergeometric test), which is derived from text-mining of the scientific literature in PubMed [49]. Moreover, miRCancer provides information about the cancer types that are associated with miRNAs in the literature; ~35% of the predictions of cancer miRNAs in specific cancer types were supported by the literature to be involved in the same cancer type (p-value = 7.02e−27; hypergeometric test).

We identified a core set of 17 miRNAs (derived from 9 pre-miRNAs) that were identified in at least five out of the seven cohorts (Figure 5A): hsa-miR-17-3p, hsa-miR-17-5p, hsa-miR-20a-3p, hsa-miR-20a-5p, hsa-miR-708-3p, hsa-miR-708-5p, hsa-miR-92a-1-5p (predicted in all 7 cohorts), hsa-miR-18a-3p, hsa-miR-18a-5p, hsa-miR-155-3p, hsa-miR-155-5p (6 cohorts), hsa-miR-205-3p, hsa-miR-205-5p, hsa-miR-324-3p, hsa-miR-324-5p, hsa-miR-629-3p, and hsa-miR-629-3p (5 cohorts). We did not observe a correlation between the number of potential target genes for a miRNA and the number of cohorts where it is predicted (Figure S20). All these miRNAs are derived from precursors of already established oncomiRs or tumor suppressor miRNAs, or known to be involved in immune response or inflammation [50–61]. Note that hsa-miR-17-3p, hsa-miR-17-5p, hsa-miR-18a-3p, hsa-miR-18a-3p, hsa-miR-20a-3p, hsa-miR-20a-5p, and hsa-miR-92a-1-5p are part of a single miRNA cluster on chromosome 13 and this polycistronic cluster (known as miR-17-92) is well known to be composed of oncomiRs involved in proliferation and tumor angiogenesis, and reducing apoptosis of cancer cells [50].

**Figure 5.**
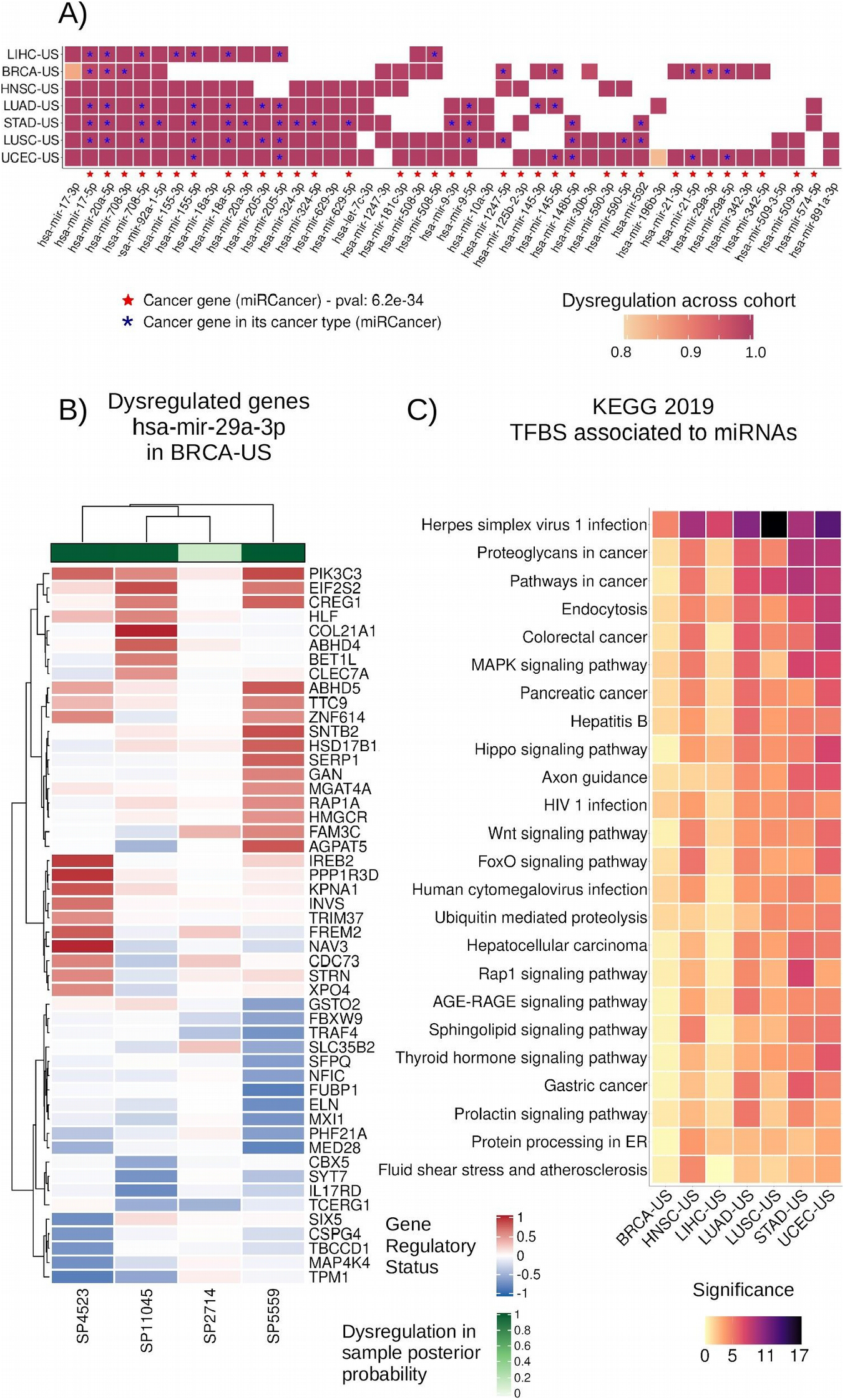
Overview of miRNA driver predictions and their dysregulated target networks. **A)** miRNAs predicted as potential drivers by *xseq* in at least two TCGA cohorts. Cell colors indicate the posterior probability computed over the corresponding cohort. Red stars indicate that the miRNA is annotated as a cancer miRNA in miRCancer [48]. Blue stars indicate that the miRNA was reported as a cancer miRNA in the specific cancer type where it is predicted by *xseq*, according to miRCancer annotation. **B)** Dysregulated network of target genes for hsa-mir-29a-3p predicted in BRCA-US (rows: dysregulated targets; columns: samples with cis-regulatory mutations associated with hsa-mir-29a-3p). The color scale represents gene regulatory status posterior probability (red: up-regulation; blue: down-regulation represented as the posterior probability times −1). **C)** KEGG 2019 most enriched terms (rows) for all the dysregulated genes associated with the identified miRNA drivers across TCGA cohorts (columns). Terms are ordered by their mean rank across all cohorts. Significance is provided as −log_10_(p-value).

When visualizing the dysregulated networks of miRNA targets in samples harbouring the predicted cancer-associated miRNAs, we observed subsets of the networks as up- or down-regulated across patients from the same cohort (Figure 5B). This observation is similar to what we detected in protein-coding gene networks (Figure 4A). Note that the miRNA target networks observed with altered expression for a given miRNA may vary between cohorts for the same miRNA as some targets are specifically expressed or altered in a subset of tissues or cell types (Figure S21).

Similar to what we detected with disrupted gene networks of protein-coding genes, functional enrichment for miRNA targets with altered expression highlighted transcriptional activity terms and biological pathways associated with carcinogenesis (Figure 5C). Further, these results were recurrently found when considering disrupted target genes in each cohort independently (Figures S15-S18). We discovered several virus infection-related terms enriched across the cohorts (Figures 4B-C and 5C), arguing for a potential link between viral infections and cancer initiation/progression, as previously suggested [62,63], via miRNAs.

Altogether, this study provides a first foray in the analysis of a combined effect of transcriptional and post-transcriptional dysregulation downstream of somatic cis-regulatory mutations associated with miRNAs in cancer cells. It highlights a core set of miRNAs associated with cis-regulatory mutations that are linked to a cascading alteration of gene regulatory networks involved in cancer onset and progression.

### Complementary analysis of independent breast cancer cohorts supports several cancer-associated miRNA predictions

We aimed to assess the recurrence of the predictions for breast cancer obtained from the 92 samples of the BRCA-US cohort from TCGA in a complementary cohort. We applied the same methodology with the same parameters on the BASIS breast cancer cohort [64], which is composed of 256 breast cancer samples with the same trio of data types available (WGS, RNA-seq, and miRNA expression - from microarrays; Additional file 5).

Similar to the BRCA-US TCGA analysis on protein-coding genes, our analysis of the BASIS cohort predicted known cancer drivers identified by associating LoF or cis-regulatory mutations with dysregulation of their gene networks. Further, we observed enrichment of similar key cancer pathways when considering the dysregulated genes associated with the predicted cancer-associated genes (Figures S22-S23). Breast cancers can be categorized into estrogen receptor positive (ER+) and negative (ER−), each subtype harbouring a distinctive signature of gene expression. We explored how the distribution of ER status in patients from the two cohorts could impact the predictions of cancer-associated genes. The TCGA BRCA-US cohort is composed of approximately the same number of ER+ and ER− patients while the BASIS cohort is composed of 72% of ER+ patients. Given the size of the BASIS cohort (256 samples), it was possible to perform two additional analyses on ER+ (184 samples) and ER− samples (72 samples) independently. The analysis of cis-regulatory mutations associated with protein-coding genes revealed one prediction common to TCGA and ER+ BASIS cohorts (*IL12RB1*; Figure S24). Despite this small intersection, the functional enrichment analysis of the dysregulated genes associated to all predicted genes were similar in the two cohorts (Figures S25-S26), suggesting that although the predictions vary among cohorts with different aethiology, the dysregulated pathways are likely the same.

We predicted four miRNAs associated with cis-regulatory mutations in the BASIS cohort when considering all samples (Figure 6). Two of these miRNAs, hsa-miR-145-5p and hsa-mir-29a-3p, were previously identified by our methodology using the BRCA-US cohort. We did not predict any driver miRNAs associated with cis-regulatory mutations when examining specifically the ER+ samples. However, we identified hsa-mir-17-3p, hsa-mir-17-5p, hsa-mir-18a-5p, hsa-mir-20a-5p, and hsa-mir-155-5p when considering ER− samples. These five miRNAs were recurrently found across multiple TCGA cohorts (Figures 5–6). Note that when predicted using the BRCA-US TCGA cohort, hsa-mir-17-3p, hsa-mir-17-5p, hsa-mir-18a-5p, and hsa-mir-20a-5p miRNAs were predicted in a majority of ER− samples as well. As expected, these results confirm that the cohort clinicopathological composition impacts the predictions as it can impact the landscape gene expression distributions across samples. Nevertheless, the complementary analyses of the TCGA and BASIS breast cancer cohorts exhibited hsa-mir-145-5p, hsa-mir-29a-3p, hsa-mir-17-3p, hsa-mir-17-5p, hsa-mir-18-5p, and hsa-mir-20a-5p as recurrently predicted breast cancer-associated miRNAs linked to cis-regulatory mutations and dysregulation of their target gene networks. Functional enrichment analysis confirmed that the dysregulated miRNA target gene networks are enriched for genes involved in transcriptional regulation and in cancer-relevant pathways such as the P53 pathway, hypoxia, and DNA damage response (Figure S26).

**Figure 6.**
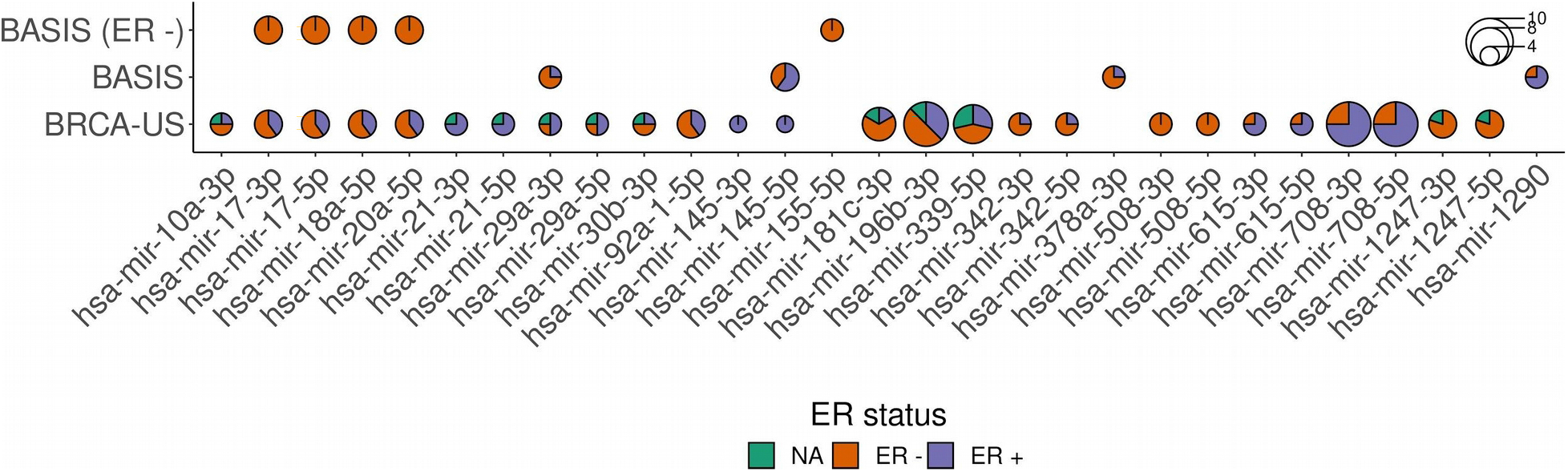
Predicted miRNA drivers in breast cancer cohorts. Predicted miRNAs in TCGA (BRCA-US) and BASIS (all samples and ER− samples only). The pie plots represent the distribution of samples with ER−/+ status.

Finally, we further evaluated the clinical potential of the predicted breast cancer miRNAs for breast cancer survival estimation. For this purpose, we considered a third cohort, METABRIC [65], which is composed of 1282 samples. We computed Kaplan-Meier survival curves and log-rank tests using miRNA expression from the METABRIC cohort for the miRNAs predicted as drivers in the BRCA-US and BASIS cohort (for 24 of the predicted miRNAs in breast cancer). Examining both overall survival and breast cancer specific survival values, we observed significant log-rank test p-values for hsa-mir-29a-3p, hsa-mir-1290, and hsa-mir-20a-5p (without multiple hypothesis correction; Figure 7 and Figures S27-S28). Note that hsa-mir-20a-5p and hsa-mir-29a-3p were recurrently predicted in our analyses of the BRCA-US and BASIS cohorts. Taken together, these results reinforce a posteriori the potential of some miRNAs we predicted as their level of expression could be used for prognosis.

**Figure 7.**
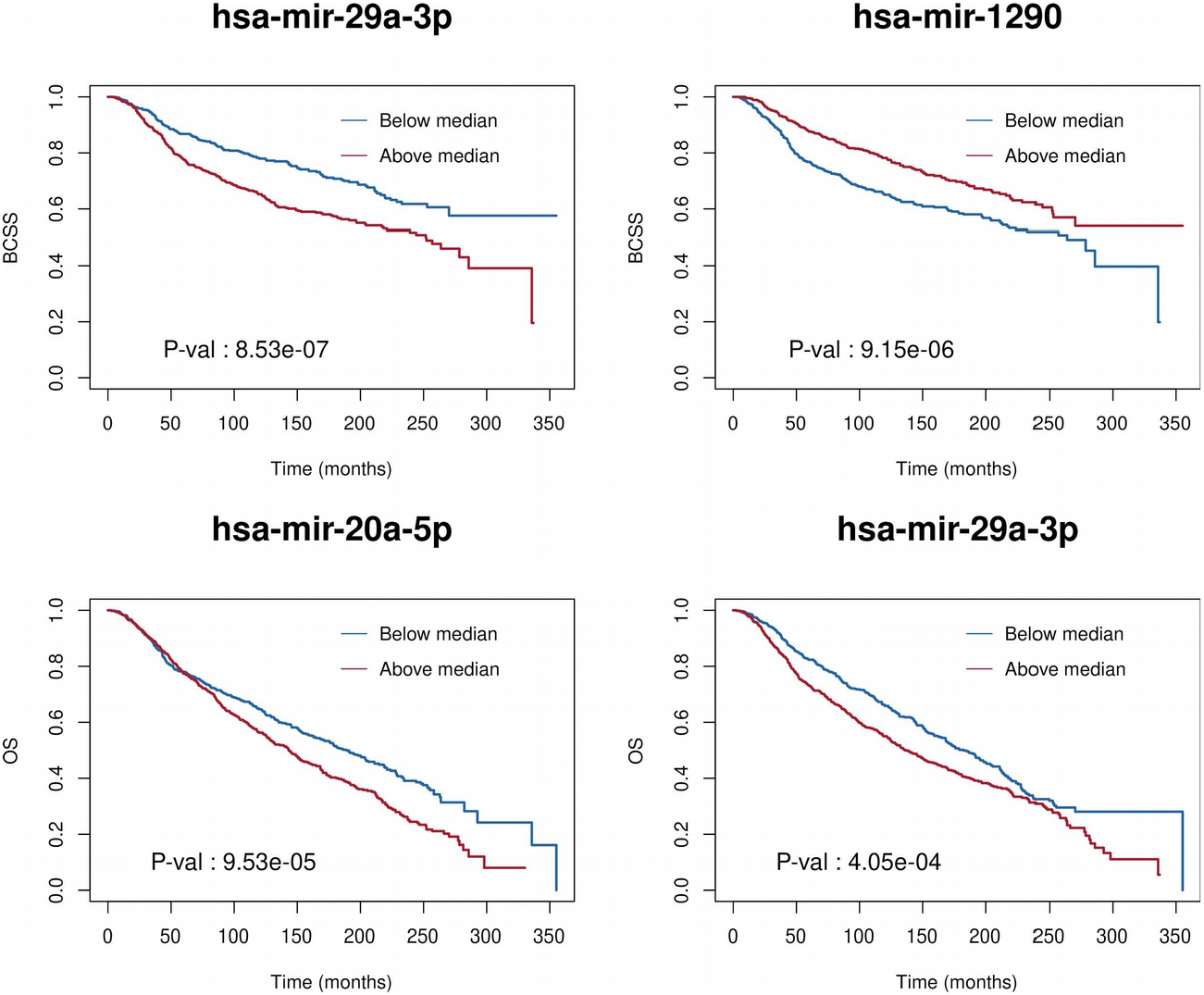
Survival curve analysis for some predicted miRNA drivers. Kaplan-Meier survival curves obtained using the METABRIC cohort for the most significant driver miRNAs identified in the breast cancer cohorts. Samples were separated into two groups according to the level of miRNA expression (above/below the median). Log-rank test p-values are indicated. OS: overall survival. BCSS: breast cancer specific survival.

## DISCUSSION

In this study, we explored how cis-regulatory mutations at TFBSs could be used to predict genes through the association of somatic mutations with a cascading trans-effect on gene regulatory network dysregulation. Contrary to most methods predicting cancer-driving events based on recurrence of mutations, we sought to couple cis-regulatory mutation information with gene expression data from the same samples to highlight direct evidence of the regulatory impact of the mutations. By integrating whole-genome somatic mutations, RNA-seq, and small RNA-seq data with gene regulatory networks, we performed pan-cancer predictions of protein-coding and miRNA genes associated with somatic cis-regulatory mutations in patients from seven distinct cancer types. Our study provides a large-scale foray to predict cancer-associated protein-coding and miRNA genes associated with somatic mutations by combining both transcriptional and post-transcriptional information. Our results provide new insights into the potential impacts and causes of the alteration of the gene regulatory program observed in cancer cells along with their cascading effects on key biological pathways.

We specifically focused on somatic mutations lying within a high-quality set of TFBSs representing direct TF-DNA interactions with both experimental and computational evidence [37] and covering ~2% of the human genome. We acknowledge that this set of TFBSs might represent a limited subset of all potential TFBSs in the human genome as it was derived from experiments available for a reduced number of TFs and cell types/tissues (231 TFs out of the ~1,600 human TFs reported [11] and 315 cell types and tissues). Moreover, some TFBSs might not be active in the cell type of origin associated with the cancer types studied here. Nevertheless, we provided evidence that the regions considered are likely enriched for functional genomic elements since they harbour mutation rates similar to what is observed in exonic regions (Figure 1). The reduced mutation rates in exons and the limited increase in surrounding regions could be attributed to increased mismatch repair and nucleotide excision repair in exons as previously shown [66,67]. Our similar observation when considering our set of TFBSs is in line with our previous observation in B-cell lymphomas [27] but somewhat in disagreement with previous studies showing that nucleotide excision repair is impaired by the binding of TFs to DNA [68,69]. We hypothesize that the differences observed could be partially explained by the fact that (1) our mutation rate analysis considered TFBSs predicted from several cell lines and tissues instead of focusing on TFs and TFBSs specific to the considered cell types or conditions (such as UV-exposure in melanoma), and (2) we did not filter TFBSs based on open chromatin data from matched cell types.

Contrary to previous studies assessing the impact of mutations on TF-DNA binding affinity or the enrichment for mutations in cis-regulatory regions [70–72], we particularly evaluated the impact of cis-regulatory mutations on expression alteration in gene networks. As such, our approach does not quantify the direct impact of mutations on the obliteration of TF-DNA interactions but uses RNA information as the ultimate readout. A previous method systematically assessed the potential impact of somatic mutations in genomic tiles near genes’ TSSs on gene expression [25]. Here, we considered mutations lying within a specific set of TFBSs without restrictions on distances to TSSs and evaluated the trans-association of the mutations with genes’ network deregulation. Nevertheless, we acknowledge that the method misses mutations that would create new TFBSs as it is restricted to a set of predefined TFBSs.

The analysis of protein coding genes showed 31 genes that were predicted in at least two (out of the seven) TCGA cohorts analyzed, with many already known cancer drivers (Figure 2A). We observed that the predicted protein-coding driver genes through the analysis of cis-regulatory mutations did generally not contain mutations in exonic regions for the same patients (Figures 2B and S13). This observation suggests complementary mechanisms acting upon gene expression dysregulation with cascading effect on regulatory network disruption.

Given that miRNAs cover a small portion of the human genome, they harbour a small number of somatic mutations [8], limiting the possibility to affect gene expression. The potential mechanism that we propose here is the alteration of their regulatory elements. Our study highlighted cis-regulatory mutations linked to miRNAs that were associated with dysregulation of expression of the miRNA targets. In our pan-cancer analysis, we discovered a core set of 17 miRNAs associated with the dysregulation of key pathways involved in carcinogenesis. This core set of miRNAs could represent a common feature for gene expression dysregulation associated with cancer onset or progression

The analysis of the dysregulated networks associated with the predicted cancer-associated genes (protein-coding and miRNAs) shows that many genes are dysregulated in a few samples but rarely across all the mutated samples (Figure 5B). This observation suggests a phenotypic heterogeneity (i.e. alterations of different parts of the same network lead to the same phenotype). However, the functional enrichment analysis of the dysregulated genes shows consistency across cohorts and across the analyzed types of mutations (LoF and cis-regulatory) for both protein-coding and miRNA genes. Moreover, as originally described by Ding et al. [26], the *xseq* probabilistic framework can highlight the specific samples where mutations are associated with impact on gene expression, while it does not in other samples (Figure 4A). This dichotomy can in principle be used to stratify samples and mutations but, in this study, was limited by the number of samples considered.

We applied our methodology to two cohorts of breast cancer samples (TCGA BRCA-US and BASIS). Given the large number of samples in BASIS (n=256), we performed three analyses separately by considering (i) all samples, (ii) ER+ samples, and (iii) ER− samples. As expected, we observed that predictions can vary depending on the samples histopathology. This is particularly important for methods assessing impact on gene expression, which is influenced by the clinical composition of the cohorts. We acknowledge that methodological differences between the TCGA and BASIS cohorts (e.g. somatic mutation callings, small RNA-seq versus microarrays and normalization for miRNAs) could provide additional explanations for the variation in predictions. Although only one of the predicted protein-coding genes was predicted in both the BASIS and the BRCA-US cohorts, the functional enrichment analysis of the dysregulated gene networks was consistent. This observation suggests common dysregulated pathways acting as attractors that could originate from (non-recurrent) distinct cancer-associated events. It underlines the importance of addressing cancer as a disease with perturbations manifested at the gene network level. However, our miRNA analyses highlighted five miRNAs associated with cis-regulatory mutations and target gene expression alteration recurrently altered across the BRCA-US and BASIS breast cancer cohorts (Figure 6).

Despite the multiple lines of evidence for the prediction of cancer-associated genes in this study, we acknowledge that the predictions can provide false positives and false negatives due to multiple reasons such as: (i) a limited number of TFs with high-quality TFBSs; (ii) TFBS-target gene associations obtained by a naive approach combining information from an integrative database [39] and association to the closest TSS (Figure S8); we hypothesize that many of these associations may be irrelevant or incorrect and many others are missing; (iii) a diversity of tumor purity within the considered samples (despite the original threshold of 80% used by TCGA); (iv) a limited number of WGS datasets (tens of samples) within each cohort, compared to the number of samples with WXS (hundreds) used in other studies; (v) prior networks that might be incomplete. However, one of the main limitations of this project is the low number of tumor samples with both WGS and RNA-seq data, this limitation not only biases the community research towards the study of exonic regions, but also limits the statistical power of the methods assessing the impact of cis-regulatory mutations on gene network expression alteration.

Altogether, we argue that our capacity to predict cancer-associated cis-regulation mutations will increase as more high-quality TFBSs for more TFs and improved methods to associate TFBSs to their target genes become available. In addition, focusing on cis-regulatory regions specifically open or active in cancer samples would inform where somatic mutations are likely effective. We expect that with more WGS, RNA-seq, and other genomics datasets derived from cancer samples available, the community will resume the paucity in the detection of non-coding cancer-associated events [8].

## CONCLUSION

By integrating whole-genome somatic mutations, RNA-seq, and small RNA-seq data with gene regulatory networks across seven cancer types, we were able to highlight cis-regulatory mutations associated with the dysregulation of gene regulatory networks through specific protein-coding and miRNA genes. The enrichment for known cancer genes, TFs, and the functional enrichment analysis reinforce a posteriori the predicted protein-coding and miRNA genes as being involved in biological pathway alteration affecting cancer development through exonic and cis-regulatory alterations. Our study represents, to our knowledge, the first large-scale analysis of cis-regulatory mutations that are linked to gene expression alteration in key cancer-associated pathways. Our results suggest that this process can be achieved in a flexible way as we observed different genes in different patients but all associated with deregulation of the same pathways. Combining transcriptional and post-transcriptional information, we identified a core set of 17 miRNAs linked to altered cancer pathways across cancer types. These pan-cancer results here provide new insights into the impact and potential causes of miRNA-mediated gene expression dysregulation. This work extends our capacity to address the discovery gap of cancer-associated events identification through the analysis of noncoding mutations and miRNA genes.

## MATERIAL AND METHODS

All analyses were performed using the hg19 human genome assembly. When data was obtained from another human genome assembly, coordinates were converted to the hg19 assembly using the liftOver tool provided by the UCSC Genome Browser [73,74].

### Cancer patient data

We considered TCGA [38] cohort samples for which trios of (i) whole genome somatic mutations, (ii) RNA-seq, and (iii) small RNA-seq data were available with at least 30 patients per cohort. Data was downloaded from the International Cancer Genome Consortium (ICGC) portal [75] through the *icgc-get client* (Additional file 6). Altogether, we collected data for 349 samples from seven TCGA patient cohorts (35 to 92 donors per cohort; Additional file 1): BRCA-US (breast invasive carcinoma), HNSC-US (head and neck squamous cell carcinoma), LIHC-US (liver hepatocellular carcinoma), LUAD-US (lung adenocarcinoma), LUSC-US (lung squamous cell carcinoma), STAD-US (stomach adenocarcinoma), and UCEC-US (uterine corpus endometrial carcinoma).

We collected data from 256 samples from the Breast Cancer Somatic Genetics Study (BASIS) cohort [64,76] for which trios of whole genome somatic mutations, RNA-seq, and miRNA microarray data were available (Additional file 5). miRNA expression was measured using the Human miRNA Microarray Slide (Release 19.0) with Design ID 046064 (Agilent Technologies, Santa Clara, CA, USA; see ref. [64] for details).

### Somatic mutations

Somatic single nucleotide variants (SNVs) and small insertions and deletions (indels) called by the tool MuSE [77] were retrieved from the ICGC portal for TCGA samples. For BASIS samples, we retrieved SNVs and indels called by the tools CaVEMan [78] and Pindel [79], respectively, used in the original study [64].

### RNA-seq and small RNA-seq normalization

Both RNA-seq and small RNA-seq raw counts were filtered to remove all genes with 0 reads in more than 50% of the samples for a given cohort. For each cohort, both matrices (RNA-seq and small RNA-seq) of raw counts were normalized to counts per million (cpm) using the *cpm* function from the R package edgeR [80] and the cpm values were scaled by log2 conversion. To avoid zeros, we added a pseudo-count of 1. Note that small RNA-seq reads were mapped to pre-miRNA coordinates by TCGA, providing information about pre-miRNA expression and not mature miRNAs.

The normalized microarray miRNA expression matrix for BASIS samples was retrieved from the original study where normalization was performed using the 90th percentile methodology [64].

### Copy number alteration computation

We downloaded copy number alteration (CNA) values predicted using the GISTIC2 tool [81] for TCGA samples through the Firebrowse database at http://firebrowse.org (Additional file 6). BASIS CNA estimates were computed using ASCAT (v2.1.1) [82] and converted into GISTIC format with −2 for homozygous loss (nMinor+nMajor=0), −1 for hemizygous loss (nMinor+nMajor=1), 0 for normal (nMinor+nMajor=2), 1 for three copies (nMinor+nMajor=3), and 2 for more than three copies (nMinor+nMajor>3). The CNA values assigned to the protein-coding genes were used in the *xseq* analysis to remove cis-effects of CNAs on the gene expression dysregulation assessment [26].

### Mutation rate analysis

For each sample, we calculated the mutation rates by dividing the total number of mutated nucleotides within a set of regions (TFBSs, exons, and flanking regions) by the total number of nucleotides covered by the given set of regions. TFBS genomic positions were obtained from UniBind [37] (see below). Protein-coding exon coordinates were retrieved from RefSeq Curated [83] (Additional file 6). Flanking regions were computed by (i) extending TFBS or exonic regions by 100, 500, and 1000 nucleotides on both sides using the *flank* bedtools subcommand and (ii) removing regions overlapping TFBSs and exonic regions using the *subtract* bedtools subcommand. Sets of regions were independently merged using the *merge* subcommand of the bedtools [84].

Random expectation for mutation rates were computed using 150 random sets of somatic mutations and applying the mutation rate computation described above. The random sets of mutations were generated by shuffling the original coordinates within the same chromosomes using the *shuffle* subcommand of the bedtools with the -*chrom* option.

### miRNAs

Genomic coordinates of human pre-miRNAs were retrieved from miRBase v20 [45] as used to predict miRNA TSSs from CAGE data by the FANTOM5 consortium [36]. When miRNA names in the miRNA-related files (expression, survival, cancer-associated miRNAs) used in this study were mapped to older versions of miRBase (starting from version v10), we updated the names to the ones used in the latest miRBase version (v22) using the miRBaseConverter R/Bioconductor package [85].

### Transcription factor binding sites

TFBSs were retrieved from the UniBind database (2019 version) at https://unibind.uio.no [37] (Additional file 6). The TFBSs correspond to high confidence direct TF-DNA interactions with both experimental (through ChIP-seq) and computational (through position weight matrices from JASPAR [86]) evidence. TFBSs were derived from 1983 ChIP-seq experiments for 231 TFs across 315 cell types and tissues [37].

### TFBS-gene association

We used the cis-regulatory element-gene associations from the GeneHancer database (v4.9), derived from 8 sources to associate TFBSs to genes (Additional file 6; Figure S8) [39]. TFBSs overlapping a cis-regulatory element annotated in GeneHancer were associated with the corresponding gene in GeneHancer. TFBSs not overlapping annotated elements were associated with the closest TSS (for a protein-coding or a miRNA gene). We considered TSSs associated with protein-coding genes from RefSeq Curated [83] and TSSs associated with miRNAs by FANTOM5 [36]. With this approach, about half of the TFBSs were associated to protein-coding or miRNA genes using GeneHancer associations and the other half to the closest TSS.

### TFBS mutations

Somatic mutations were intersected with TFBS locations using the *intersect* subcommand of bedtools v2.25.0 [84]. All mutations in TFBSs associated with miRNAs were considered for the *xseq* analysis (see below). For mutations in TFBSs associated with protein-coding genes, we followed the approach previously used by Mathelier *et al.* for the *xseq* analysis [27]. Specifically, we restricted the analysis to mutations associated with genes potentially dysregulated in the corresponding samples. Following ref. [27], genes were considered as potentially dysregulated in a given sample in cohort *C* if its expression value *v* satisfied *v* < μ−1σ or *v* > μ+1σ where μ and σ correspond to the mean and standard deviation of the expression values of the gene in *C*.

### Loss-of-function mutations

Following Ding *et al.* [26] for protein-coding exonic regions, we considered only LoF mutations that are either (i) nonsense mutations (disruptive in-frame deletion, disruptive in-frame insertion, stop gained, start lost, stop lost, and stop retained variant), (ii) frameshift mutations (frameshift variant, initiator codon variant), or (iii) splice-site mutations (splice region variant, splice donor variant, splice acceptor variant). The analysis was performed using somatic mutation data obtained from whole exon sequencing in the same TCGA samples as for the other analyses.

### Gene networks

Protein-coding gene networks were retrieved from ref. [26] and were composed of 898,032 interactions. Briefly, the networks were constructed by combining gene associations from STRING v9.1 functional protein association [87], KEGG pathway datasets [40], WikiPathway [41], and BioCyc [88] as integrated in the IntPath database [89], and TF-target links from ENCODE [90] (see ref. [26] for more details). We updated the weights of the connections whenever possible using the methods provided in *xseq*, following the methodology described in ref. [26]. Specifically, the weight between a given gene *g* and a biological partner gene *p* was set to 1 if *p* was found differentially expressed (Benjamini-Hochberg adjusted p-value ≤ 0.05) in samples where *g* is mutated in the same cohort (see Material and methods in ref. [26] for details). If there existed such genes *p*, then only these genes were kept connected to *g*. Original weights were kept otherwise.

### miRNA-target networks

miRNAs were associated with potential target protein-coding genes using predictions from TargetScan v7.2 [31]. From the list of targets for each miRNA, we filtered out the targets with less than two predicted binding sites for the given miRNA to reduce false positives [46,47]. miRNA-target weights were computed as *t_score* / 100 where *t_score* corresponds to the *targetScan context*++ score percentiles from TargetScan. We updated the weights of the connections whenever possible following the same strategy as for protein-coding genes (see above).

### *xseq* analyses

The likely associations between mutations and dysregulation of gene or miRNA target networks were calculated with *xseq* [26]. This method requires as input: a gene expression matrix of samples (RNA-seq matrix), a binary sample-gene mutation matrix, and a weighted network of connected genes. It outputs posterior probabilities associated to: (i) a sample-specific gene regulatory status (GRS, the probability of a given gene being dysregulated in a sample) for each gene connected to the gene associated with a mutation in a given sample, (ii) a sample-specific dysregulation probability (SSD, the probability that a mutation in a given gene in a given sample is associated with dysregulation of the gene’s network), and (iii) a dysregulation across the cohort probability (DAC, the probability that mutations in a gene are associated with the dysregulation of its network across patients) (Figure S9). In a first step, we removed lowly expressed genes in a cohort following the approach described by Ding et al. [26]. Briefly, *xseq* considers the 90th percentile of expression for each gene and decomposes the distribution of these values into two Gaussian distributions corresponding to low and high expression values. We considered for further analysis the genes for which their 90th percentile of expression values lie within the high expression distribution with a posterior probability ≥ 0.8 (see Ding et al. [26] for details). Next, *xseq* was used to compute all the posterior probabilities to predict genes and cis-regulatory mutations in the cancer patient cohorts. We considered potential cancer-associated genes the ones with DAC ≥ 0.8 and SSD ≥ 0.5 in at least two samples.

### Dysregulation heatmaps

The dysregulated networks for predicted protein-coding and miRNA genes are visualized as heatmaps where columns correspond to mutated samples and rows to connected genes. Heatmaps were constructed with connected genes dysregulated (GRS ≥ 0.5) in at least one sample with SSD ≥ 0.5. These genes are referred to as dysregulated genes.

### Aggregated and sample-specific networks

To evaluate whether the protein-coding genes predicted by cis-regulatory mutations are connected in the filtered networks (see Gene networks section), we built an aggregate network using all the predicted protein-coding genes within a cohort. We counted the number of disconnected subgraphs using the R packages *igraph* [91] and *ggnetwork* [92]. Similarly, we built sample-specific networks and counted the number of subgraphs in each sample, only considering the predicted genes with DAC ≥ 0.8 and SSD ≥ 0.5.

### Functional enrichment analysis

Given a list of dysregulated genes, functional enrichment analyses were performed using the R package enrichR [44] for the following databases: KEGG_2019_Human, WikiPathways_2019_Human, GO_Biological_Process_2018, and Panther_2016.

### Enrichment for cancer genes and TFs

Given a set of genes, we assessed their enrichment for cancer genes or TFs using hypergeometric tests using the *stats::phyper* function in R. The list of cancer protein-coding genes considered was constructed by considering genes that appear in at least two of the following databases: Network Cancer Gene [93], inToGen [94], and Cancer Gene Census [95]. Cancer miRNA genes were retrieved from miRCancer [48] (with data from May 1st 2019). TF genes were retrieved from the human transcription factor database [11].

### Survival analysis

To test whether miRNA expression was associated with survival, we used the METABRIC breast cancer cohort [65] with miRNA microarray expression [96] available for 1282 tumors. Expression values were downloaded from the European Genome-Phenome Archive, www.ebi.ac.uk/ega, accession number EGAS00000000122. Follow-up data were available from Curtis *et al.* [65]. Kaplan-Meier survival analyses and log-rank tests were performed using the R package *survival* with tumors separated into “high” or “low” miRNA expression groups depending on expression values above or below the median.

### Results accessibility

The analysis with all the scripts and parameters can be found through the following link: https://bitbucket.org/CBGR/workspace/projects/DYS. We provide (i) the source code for the analysis at https://bitbucket.org/CBGR/dysmir_manuscript/src/master/ and (ii) a pipeline for users to run similar analysis with their own data at https://bitbucket.org/CBGR/dysmir_pipeline/src/master/.

## SUPPLEMENTARY DATA

Additional file 1. **TCGA selected samples (specimen) with their ICGC/TCGA IDs**. Samples were selected to provide WGS, RNA-seq, small RNA-seq, and CNA data. All samples considered in this study correspond to solid tumors.

Additional file 2. **Summary of the number of mutations in the TCGA cohorts**. Basic summary statistics (median, mean, and standard deviation) of the total number of mutations and the number of mutations overlapping TFBSs.

Additional file 3. **Identified driver genes (miRNAs and protein coding genes) with their dysregulated targets**.

Additional file 4. **Summary of number of identified drivers and the number of dysregulated genes per cohort**.

Additional file 5. **BASIS selected samples and their ER status**. Samples were selected to provide WGS, RNA-seq, miRNA microarray expression, and CNA data.

Additional file 6. **Description of the resources used for this manuscript**. Brief description of the resources (gene lists, databases, external files) used to perform this study, with links to their corresponding sites and the date when they were downloaded.

## ACKNOWLEDGMENTS

As research parasites [97], we thank the ICGC and BASIS consortia and other researchers for making their data publicly available. We thank Jiarui Ding for his help on using xseq for this study; Marcel Smid for providing raw RNA-seq data for the BASIS cohort; Georgios Magklaras and Georgios Marselis for their IT support; Ingrid Kjelsvik and Elisa Bjørgo for administrative support; Rafael Riudavets Puig, Roza Berhanu Lemma, Sebastian Waszak, Marieke Kuijjer, and Yuvia A. Perez-Rico for helpful comments on the manuscript; and the Mathelier and Kristensen’s groups for insightful discussions throughout the course of execution of this project.

## FUNDING

Norwegian Research Council [187615]; Helse Sør-Øst; University of Oslo through the Centre for Molecular Medicine Norway (NCMM) (to AM and JACM); Norwegian Research Council [288404 to JACM and Mathelier group]; The Norwegian Cancer Society [197884 to Mathelier group]. MRA was a postdoctoral fellow of the South Eastern Norway Health Authority (grant 2014021 to ALBD) and a research fellow of the Norwegian Cancer Society (grant 711164 to VNK).

## Conflict of interest statement

None declared.

## CONTRIBUTION

AM was responsible for the project conception and supervision. JACM was responsible for the analysis design and execution, and for its implementation. JACM and MRA undertook bioinformatic analysis. JACM, MRA, and AM wrote the manuscript with input from all co-authors. OCL contributed with CNA values for the BASIS cohort. JWMM contributed with clinical data. AL managed samples and clinical data. AM, JACM, MRA, ALBD, and VK contributed to the data analysis and scientific input. All authors read and approved the final manuscript.

## Notes

### Competing Interest Statement

The authors have declared no competing interest.

