## Supplementary Figures for "Cis-regulatory mutations associate with transcriptional and post-transcriptional deregulation of the gene regulatory program in cancers"

Supplemental Material

Jaime A. Castro-Mondragon, Miriam Ragle Aure, Ole Christian Lingjærde, Anita Langerød, John W. M. Martens, Anne-Lise Børresen-Dale, Vessela Kristensen, and Anthony Mathelier

Last update: 2020-06-23

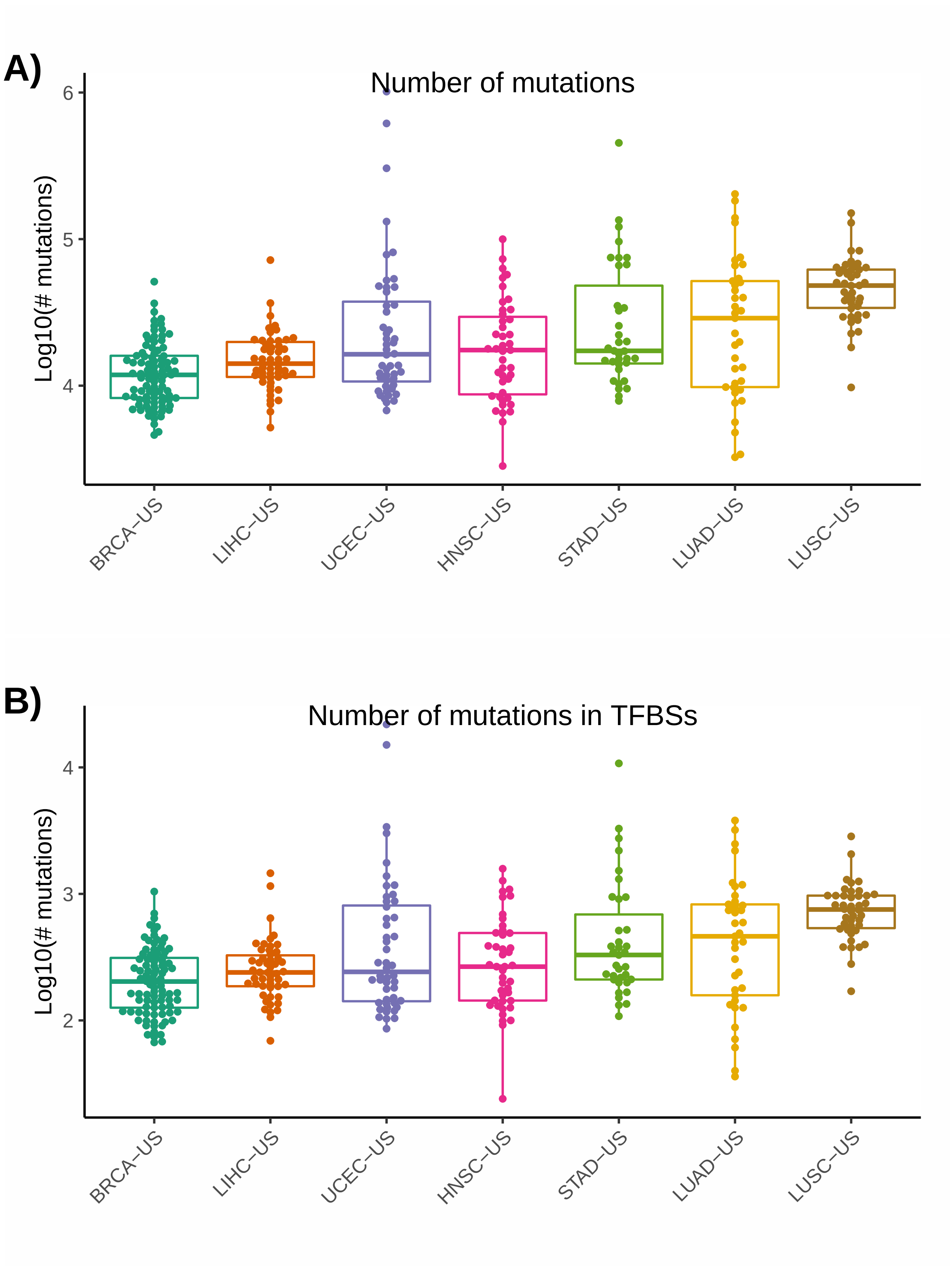


Figure S1: A) Number of mutations per sample per cohort. B) Number of cis-regulatory mutations (i.e. within TFBSs) per sample per cohort.

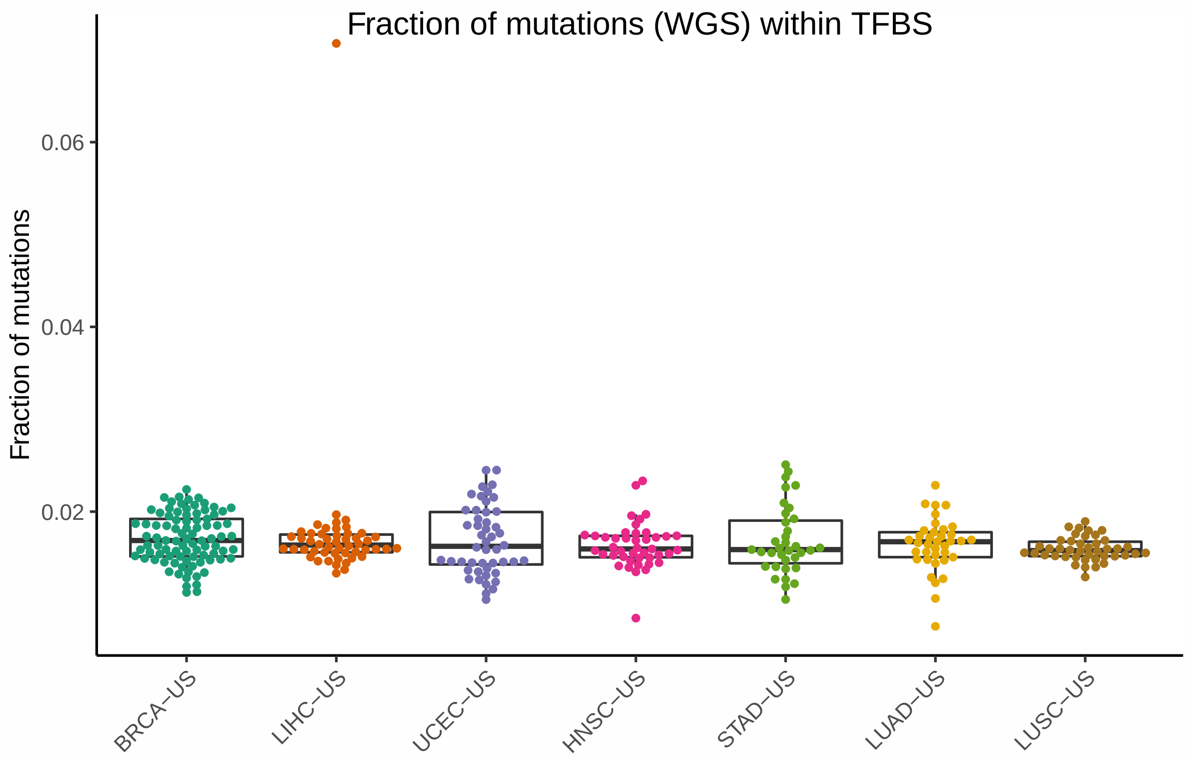


Figure S2: Fraction of somatic mutations per sample per cohort categorized as cis-regulatory mutations (i.e. within TFBSs).

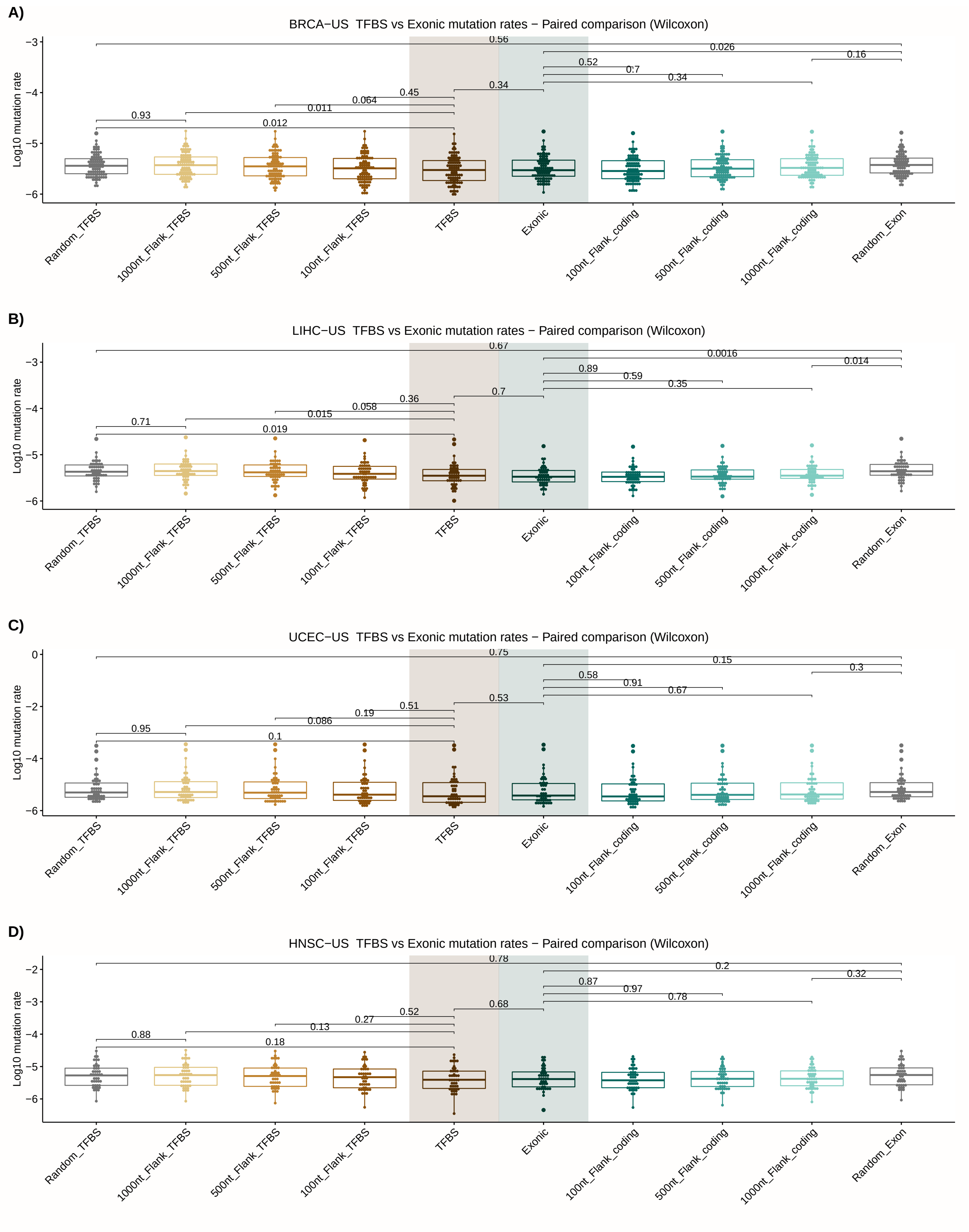


Figure S3: Paired comparisons (wilcoxon tests) of mutation rates in A) BRCA samples (n = 92), B) LIHC (n = 50), C) UCEC (n = 48), and D) HNSC (n = 43) at TFBS, exonic, flanking, and random regions.

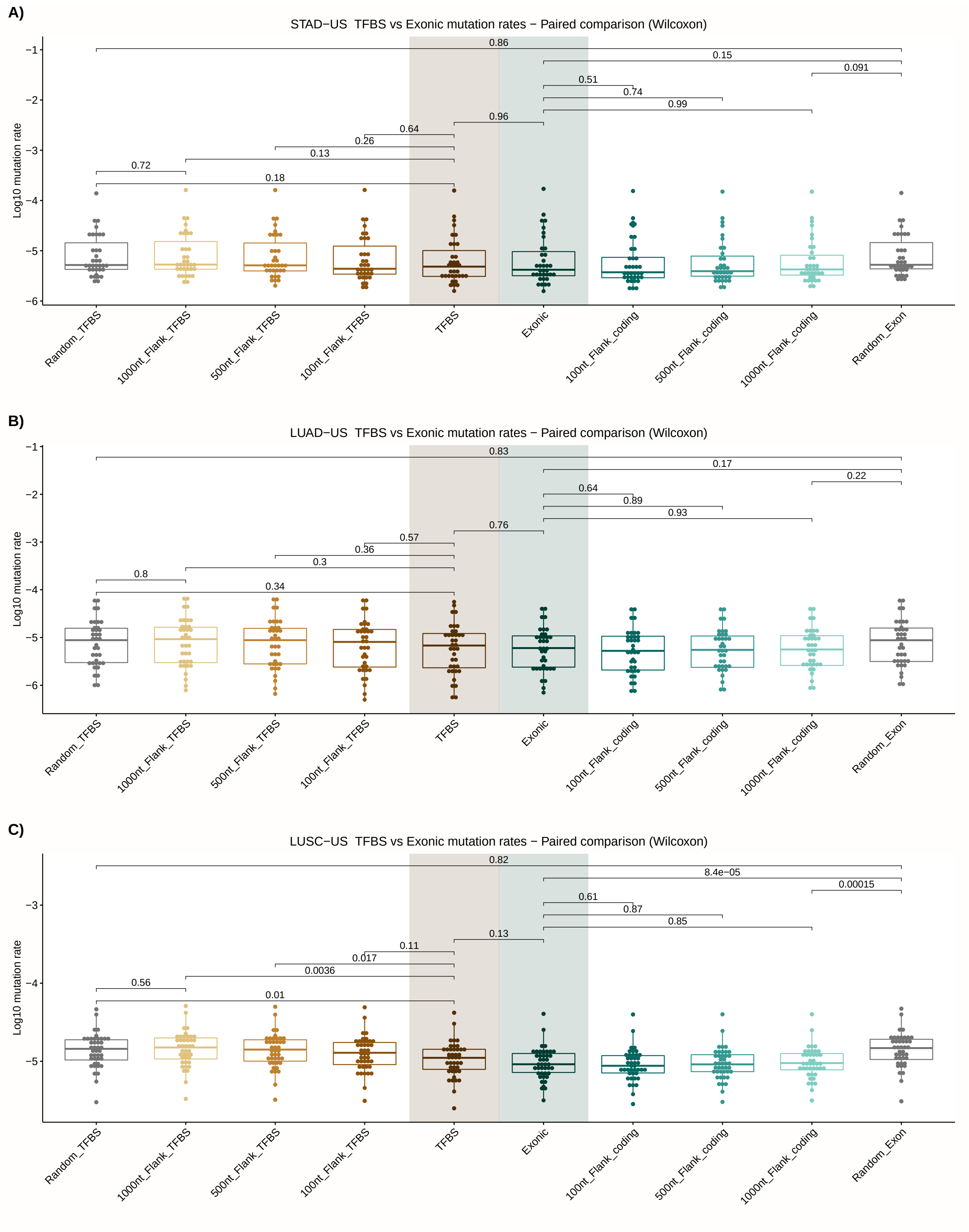


Figure S4: Paired comparisons (wilcoxon tests) of mutation rates in A) STAD samples (n = 35), B) LUAD (n = 37), and D) LUSC (n = 42) at TFBS, exonic, flanking, and random regions.

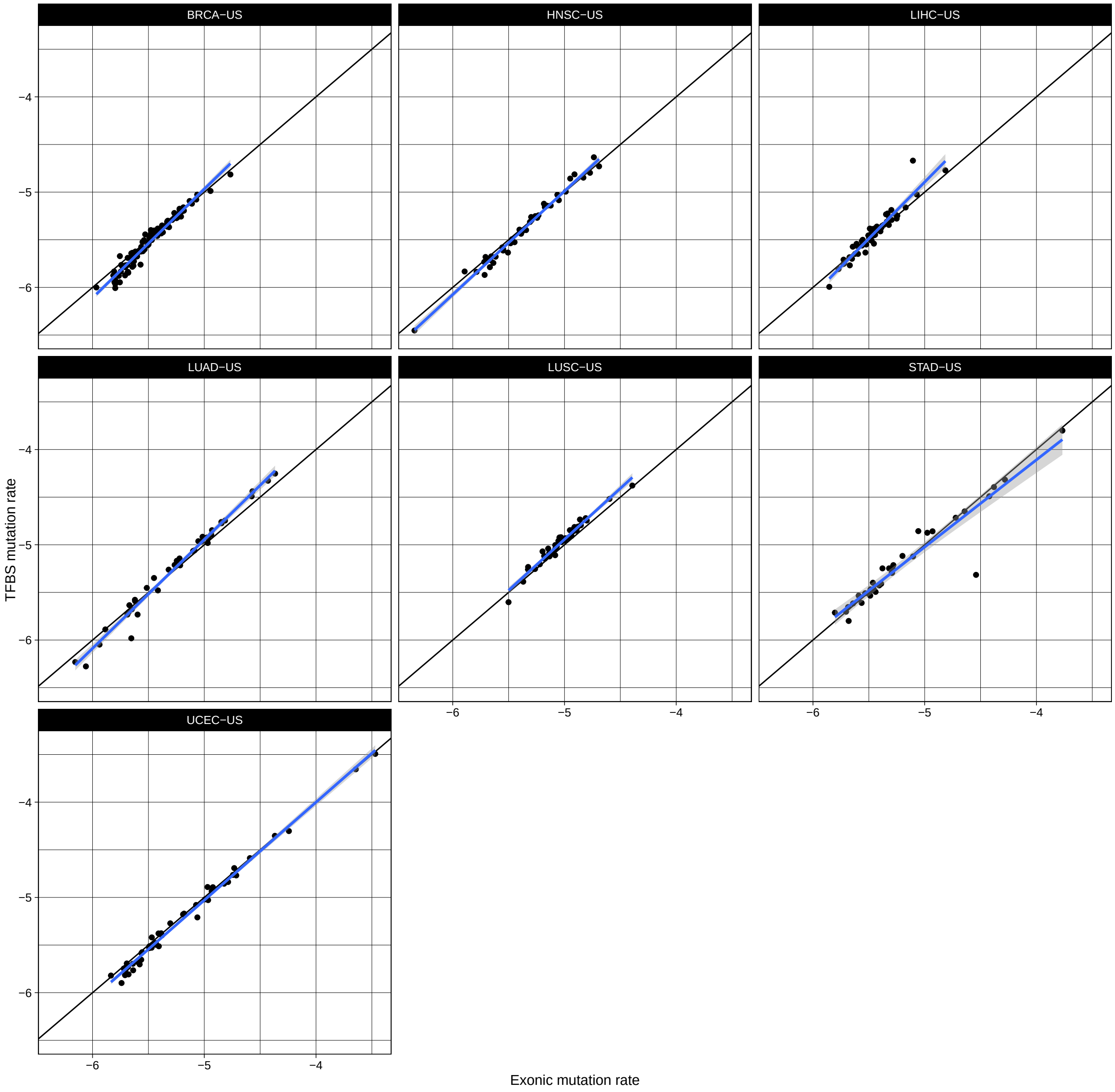


Figure S5: TFBS versus exonic mutation rates in TCGA cohorts. Both axis are -$\log_{10}$ converted. The diagonal line indicates identical mutation rates in TFBS and exonic regions. The blue line represents a linear regression model best fitting the points with a 95% confidence interval (grey zone).

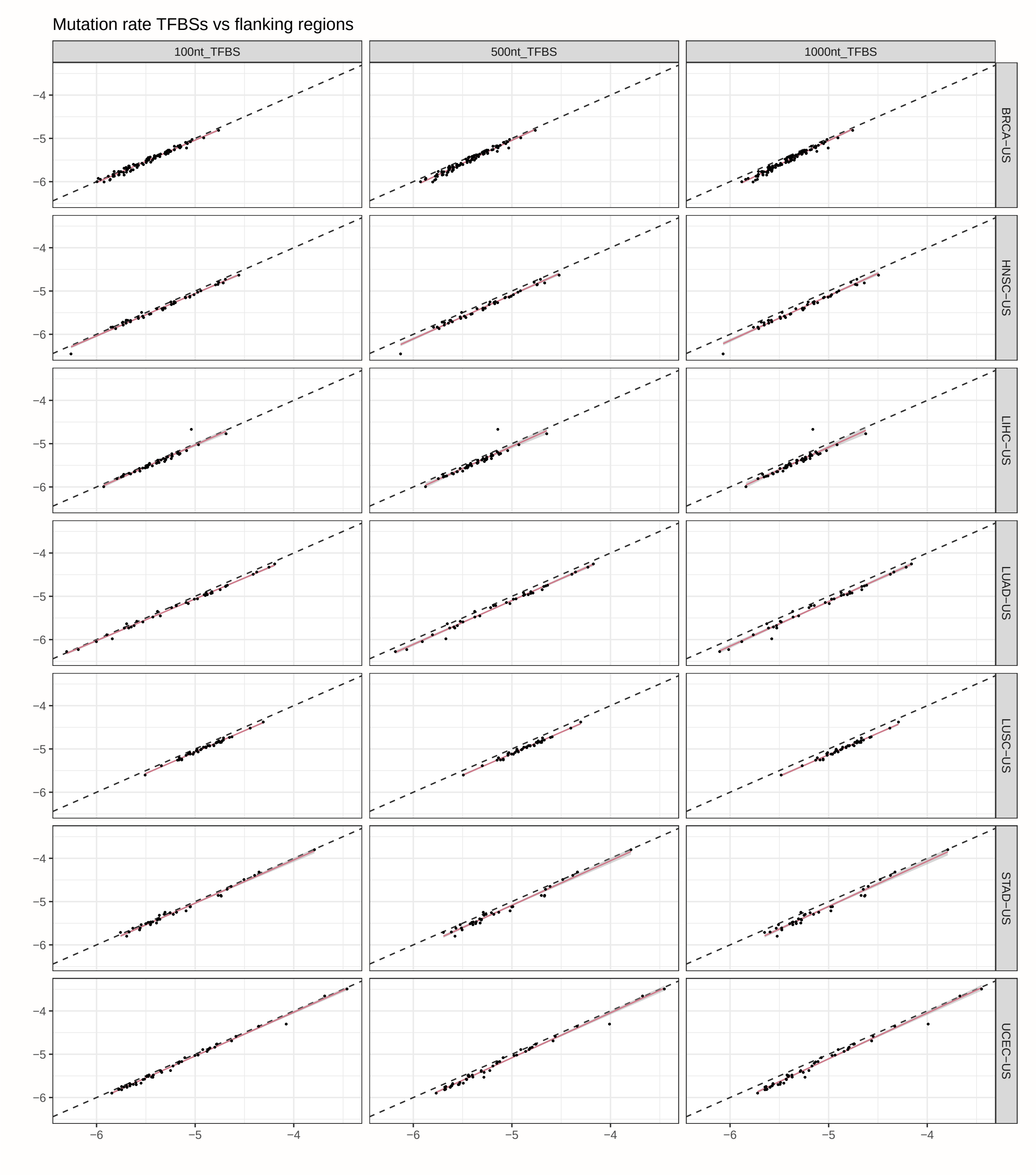


Figure S6: Mutation rates in TFBSs and their flanking regions. Left: ±50 bp, center: ±250 bp, right: ±500 bp.

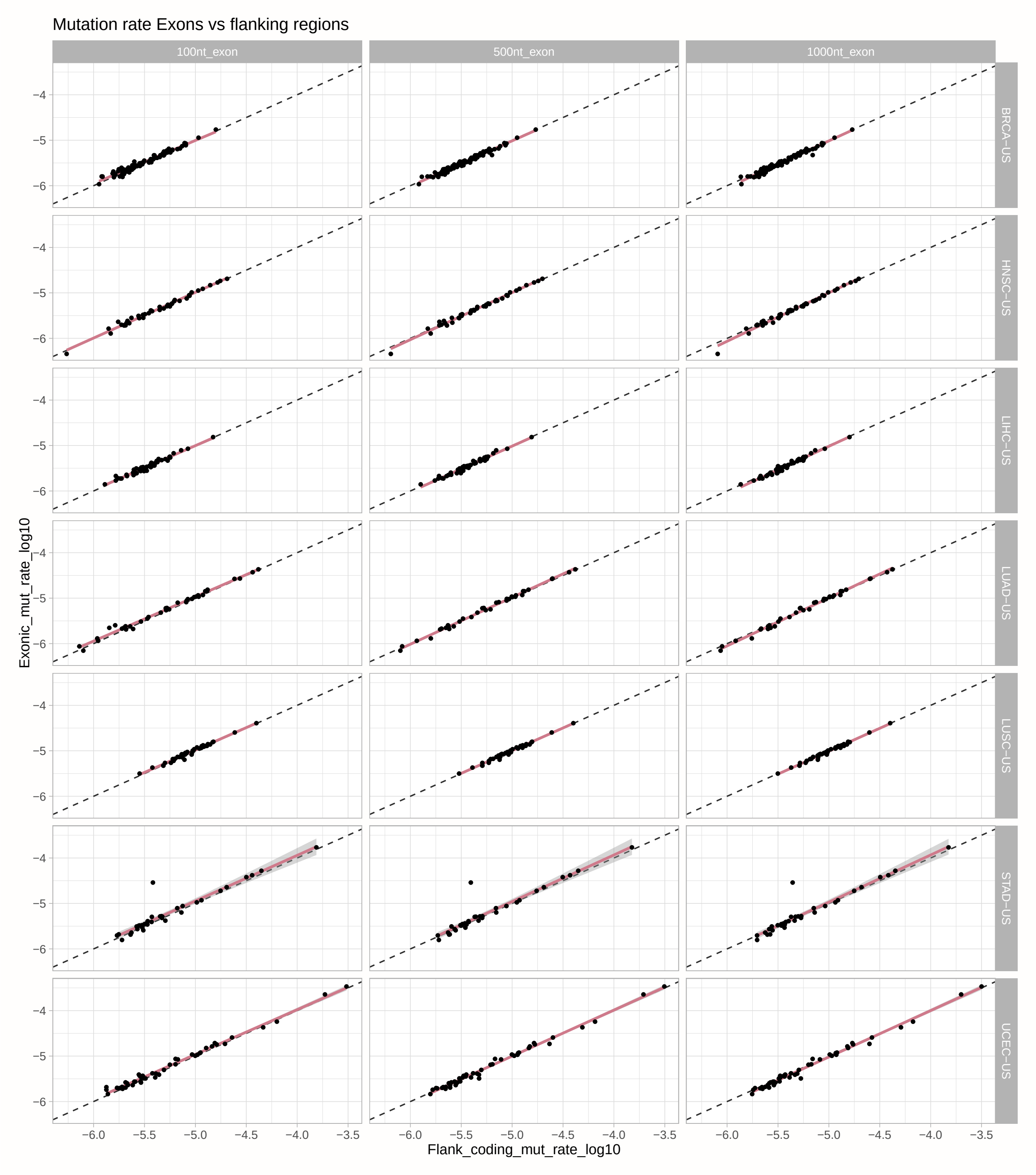


Figure S7: Mutation rates in exons and their flanking regions. Left: ±50 bp, center: ±250 bp, right: ±500 bp.

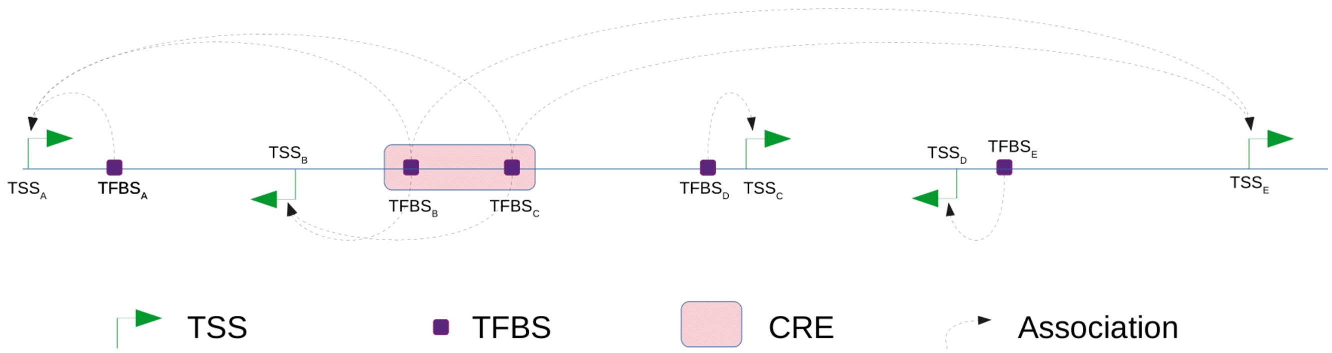


Figure S8: TFBS-gene associations. When a TFBS (purple box) lies within an GeneHancer-annotated cis-regulatory element (CRE; pink box), the TFBS is linked to its target genes according to GeneHancer, (see TFBS$_B$ and TFBS$_C$). When a TFBS is outside an annotated CRE (see TFBS$_A$, TFBS$_D$, and TFBS$_E$), it is associated with the closest TSS.

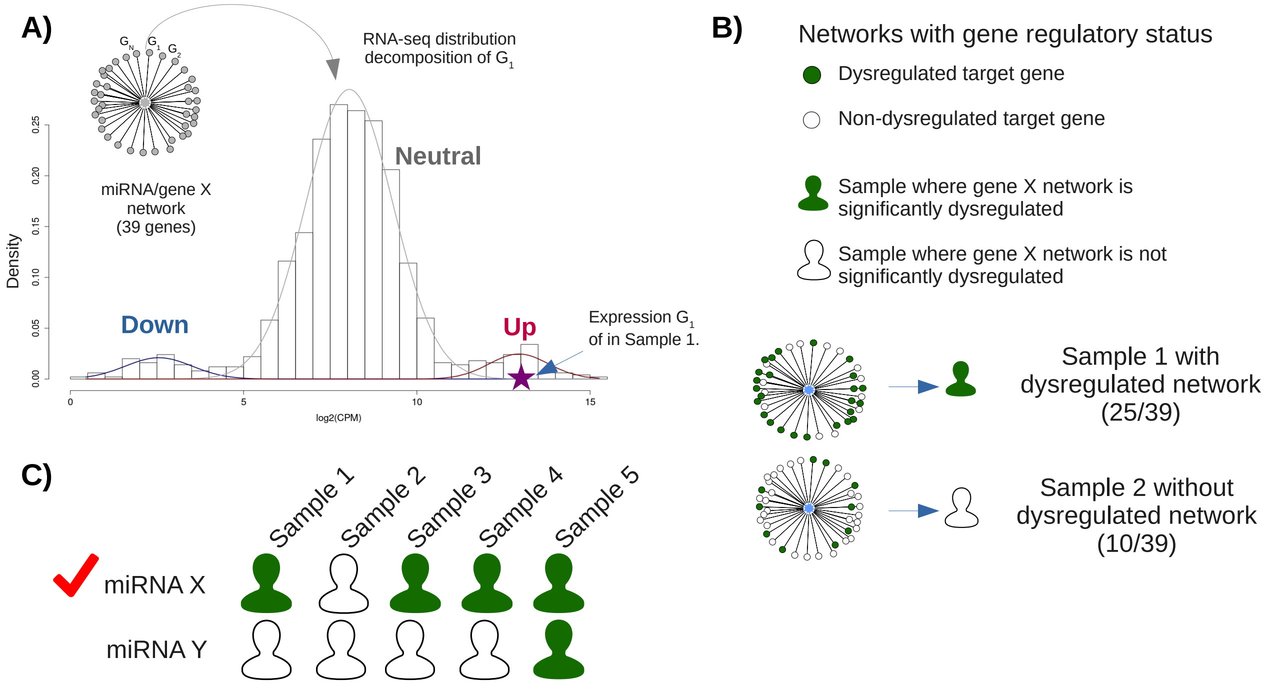


Figure S9: Likelihood association of mutations with gene expression dysregulation in xseq. Example with a network of target genes for a given miRNA. A) Gene regulatory status. For each target $i$ connected to the miRNA gene $X$ associated with a cis-regulatory mutation, xseq decomposes the expression distribution of $i$ (using all samples in the cohort) as a 3-component mixture model (down, neutral, and up). Next, xseq computes the posterior probability of observing the expression of $i$ in the mutated sample $S$ as belonging to the down, neutral, and up components. The component with the highest posterior probability provides information about the regulatory status of $i$ (down, neutral, or up). This step is repeated for each gene in the network and for each sample in the cohort. B) The sample-specific network dysregulation probability is calculated from the down and up regulation posterior probabilities of all the genes in the network for a given samples where the mutation was observed. C) The posterior probabilities of sample-specific dysregulation are used to compute the posterior probability of association between a cis-regulatory mutated miRNA and its target network dysregulation across a cohort. For a detailed explanation of the parameters and the graphical model, see the xseq publication by Ding et al., Nature Communications, 2015.

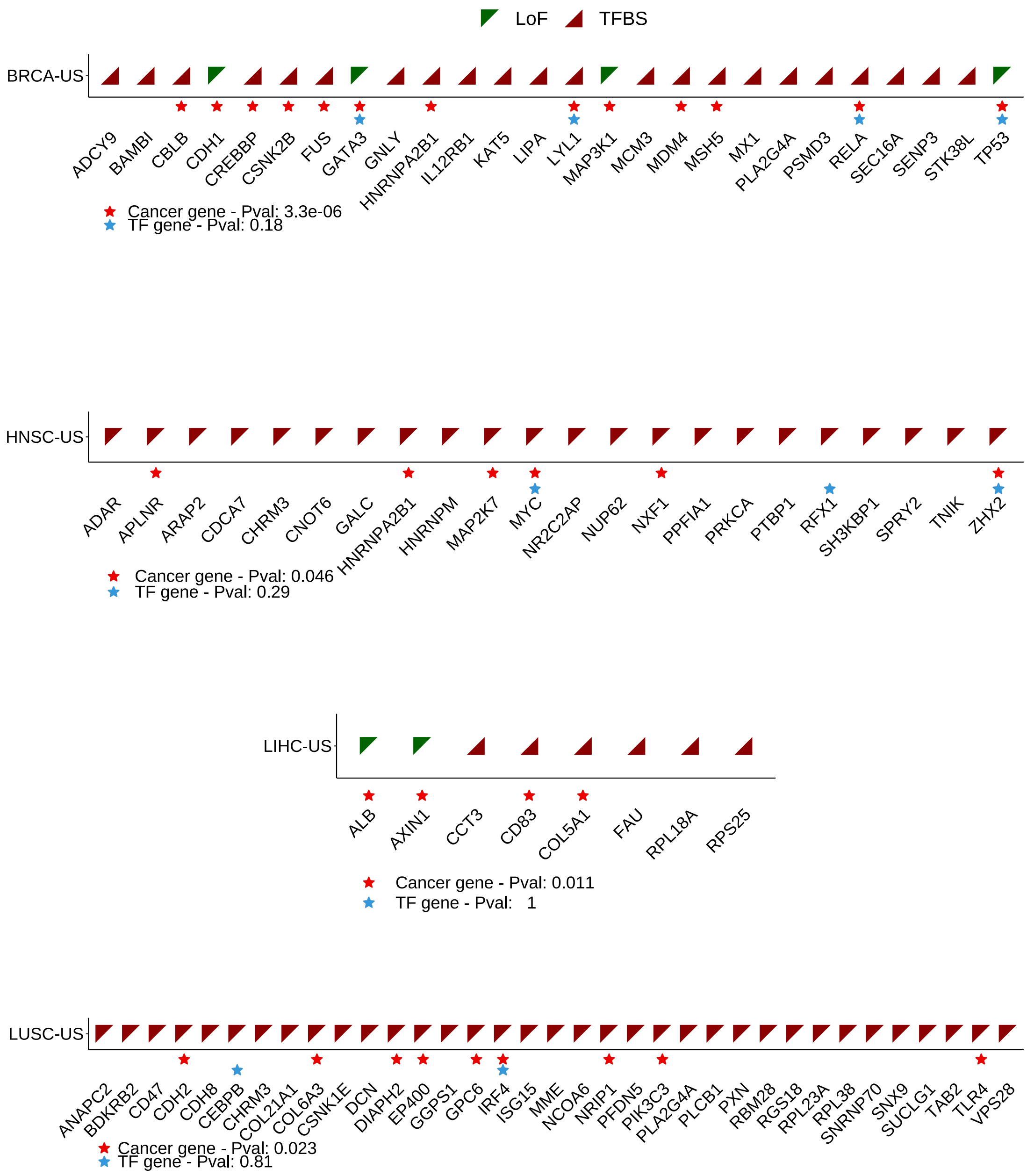


Figure S10: Protein-coding genes predicted in BRCA-US, HNSC-US, LIHC-US, and LUSC-US TCGA cohorts. Predictions were obtained applying the xseq tool when considering protein-coding genes mutated through either LoF (red triangles) or cis-regulatory (TFBS; green triangles) mutations, independently. Genes known as cancer genes (red stars) and TFs (blue stars) are highlighted and hypergeometric tests p-values for enrichment are provided in the legend (Material and methods).

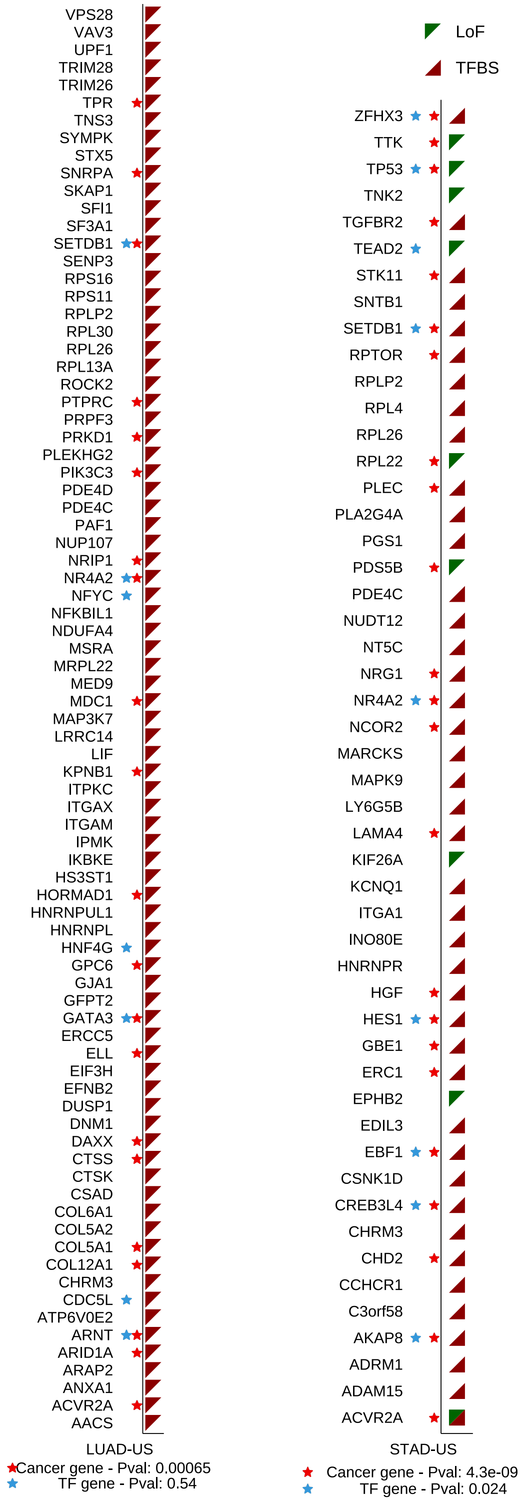


Figure S11: Protein-coding genes predicted in LUAD-US and STAD-US TCGA cohorts. Predictions were obtained applying the xseq tool when considering protein-coding genes mutated through either LoF (red triangles) or cis-regulatory (TFBS; green triangles) mutations, independently. Genes known as cancer genes (red stars) and TFs (blue stars) are highlighted and hypergeometric tests p-values for enrichment are provided in the legend (Material and methods).

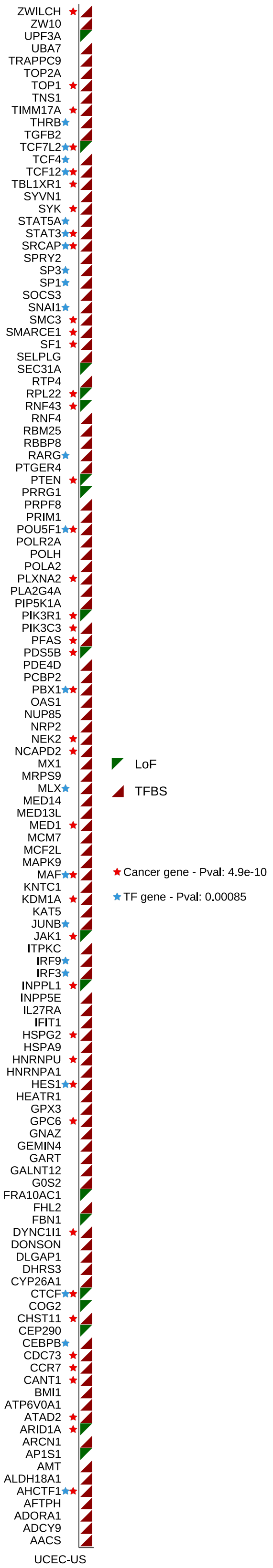


Figure S12: Protein-coding genes predicted in the UCEC-US TCGA cohort. Predictions were obtained applying the xseq tool when considering protein-coding genes mutated through either LoF (red triangles) or cis-regulatory (TFBS; green triangles) mutations, independently. Genes known as cancer genes (red stars) and TFs (blue stars) are highlighted and hypergeometric tests p-values for enrichment are provided in the legend (Material and methods).

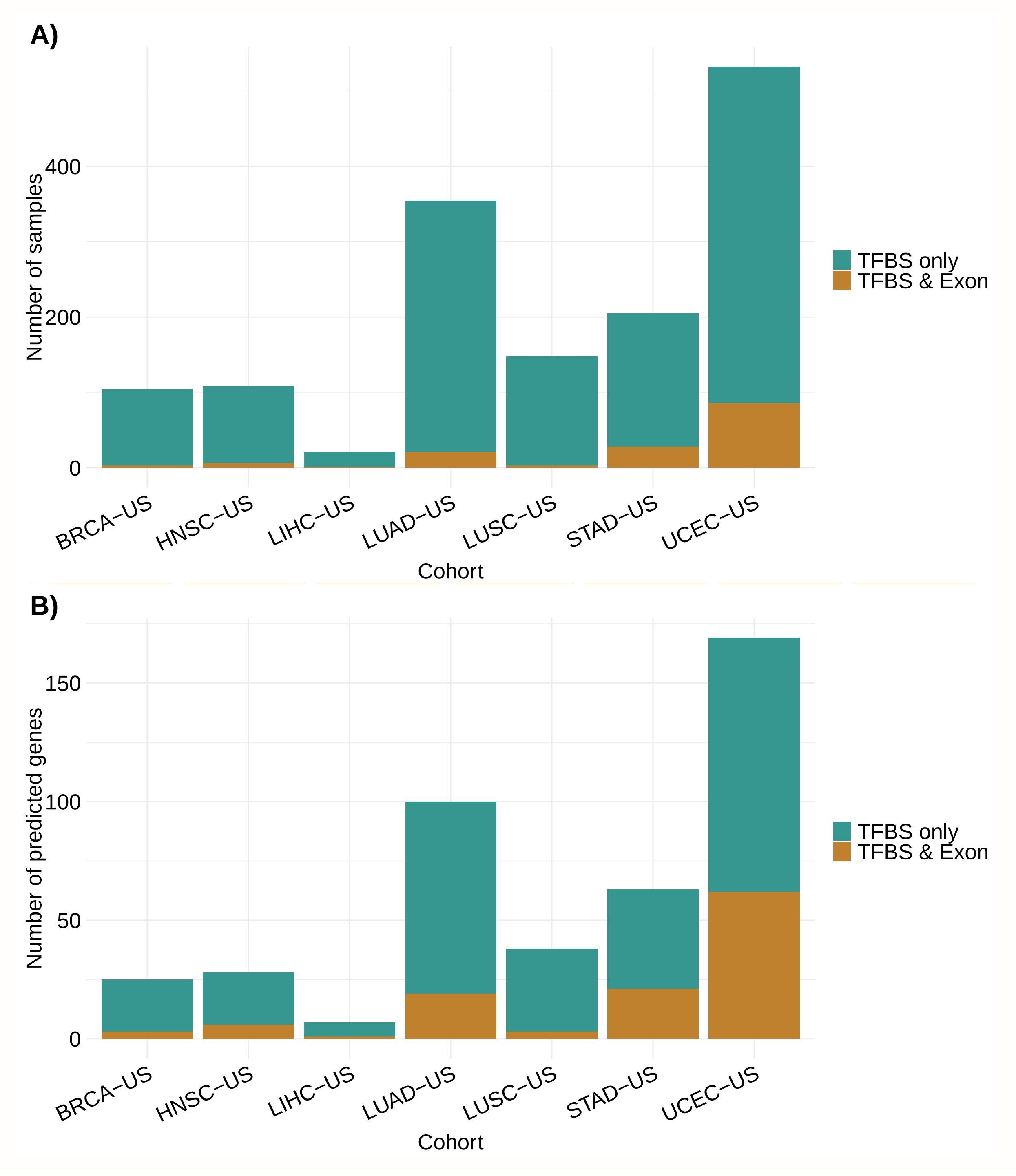


Figure S13: Protein-coding genes predicted through cis-regulatory mutations (i.e. within TFBSs) are generally not mutated in their exons in the same patients. A) For each cohort, number of samples where the predicted protein-coding genes through cis-regulatory mutations (TFBSs) also contain a LoF mutation. B) Same as in A) but providing the corresponding number of predicted protein-coding genes per cohort for each category.

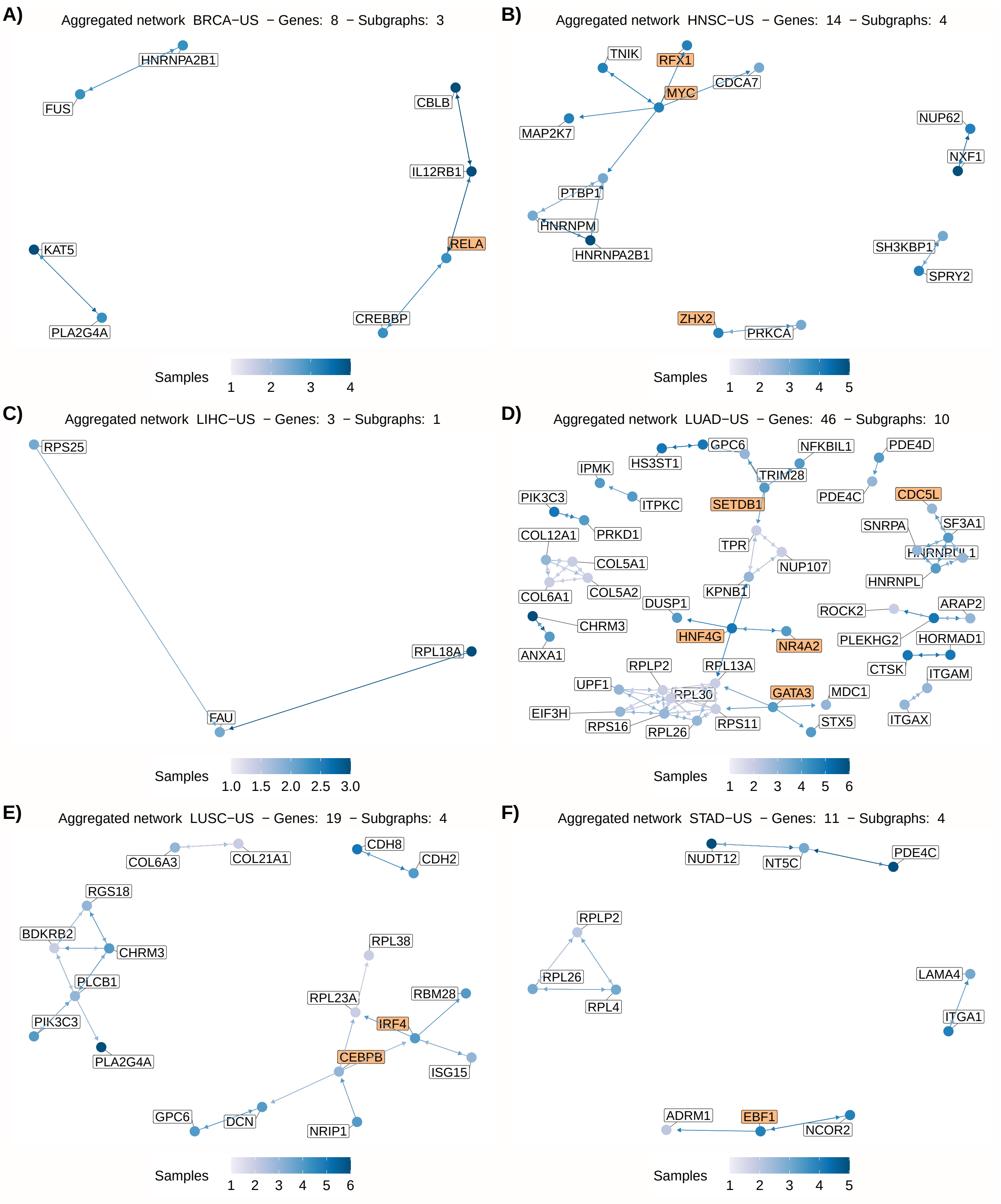


Figure S14: Network of all predicted genes in A) BRCA, B) HNSC, C) LIHC, D) LUAD, E) LUSC, and F) STAD cohorts. The node colors indicate the number of samples where the gene was predicted. TFs genes are highlighted with orange background.

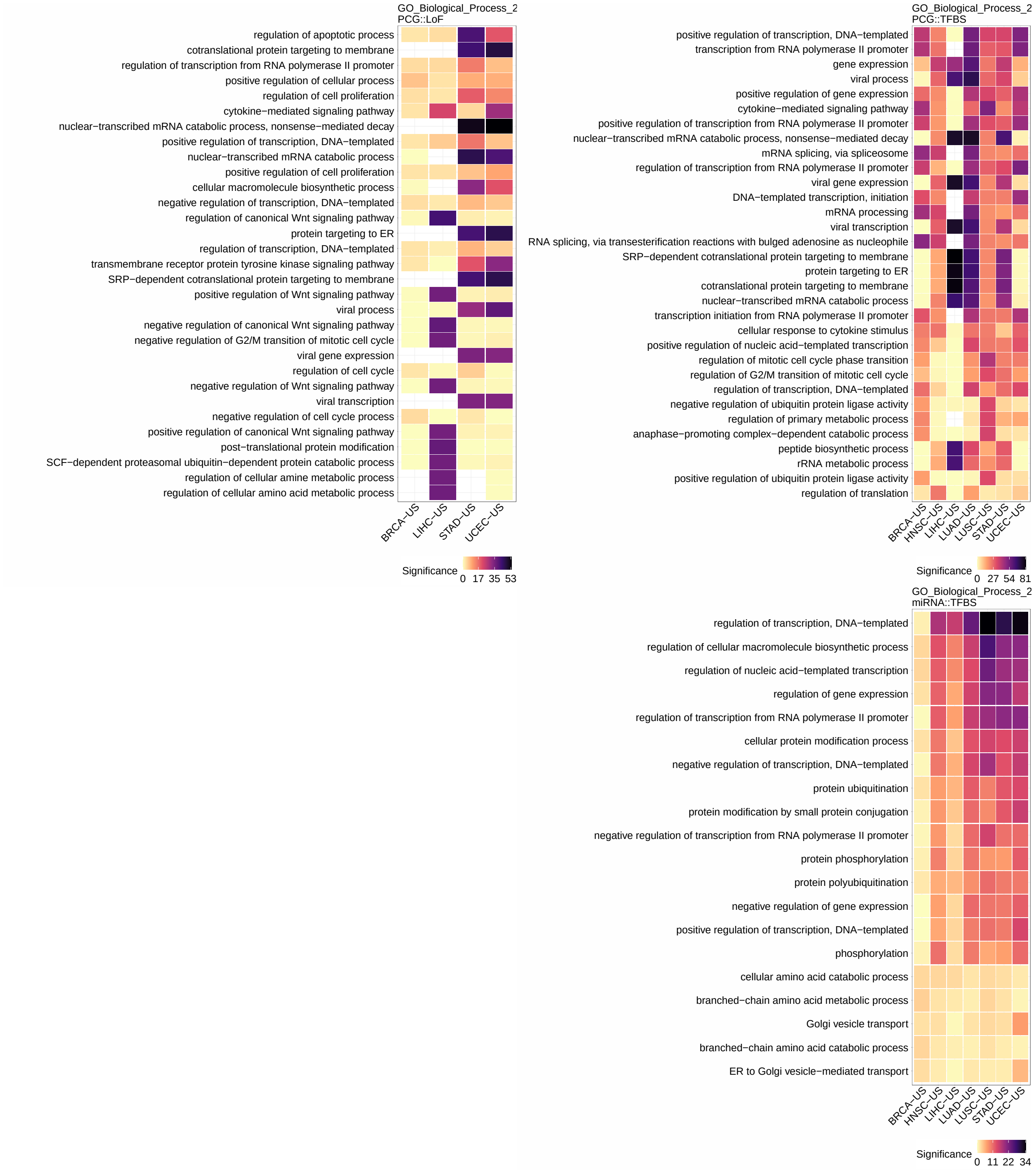


Figure S15: Functional enrichment (GO Biological Process 2018) analysis considering dysregulated genes in the networks of predicted miRNA and protein-coding genes. The heatmaps represent the significance (-$\log_{10}$(P-value)) of the most enriched terms (rows) associated to the dysregulated genes found on the seven analyzed cohorts (columns) when considering protein-coding genes with LoF mutations (PCGs::LoF; top-left), protein-coding genes with cis-regulatory mutations (PCG::TFBS; top-right), and miRNAs with cis-regulatory mutations (miRNAs::TFBS; bottom-right). The terms are ordered by their mean ranks across the cohorts.

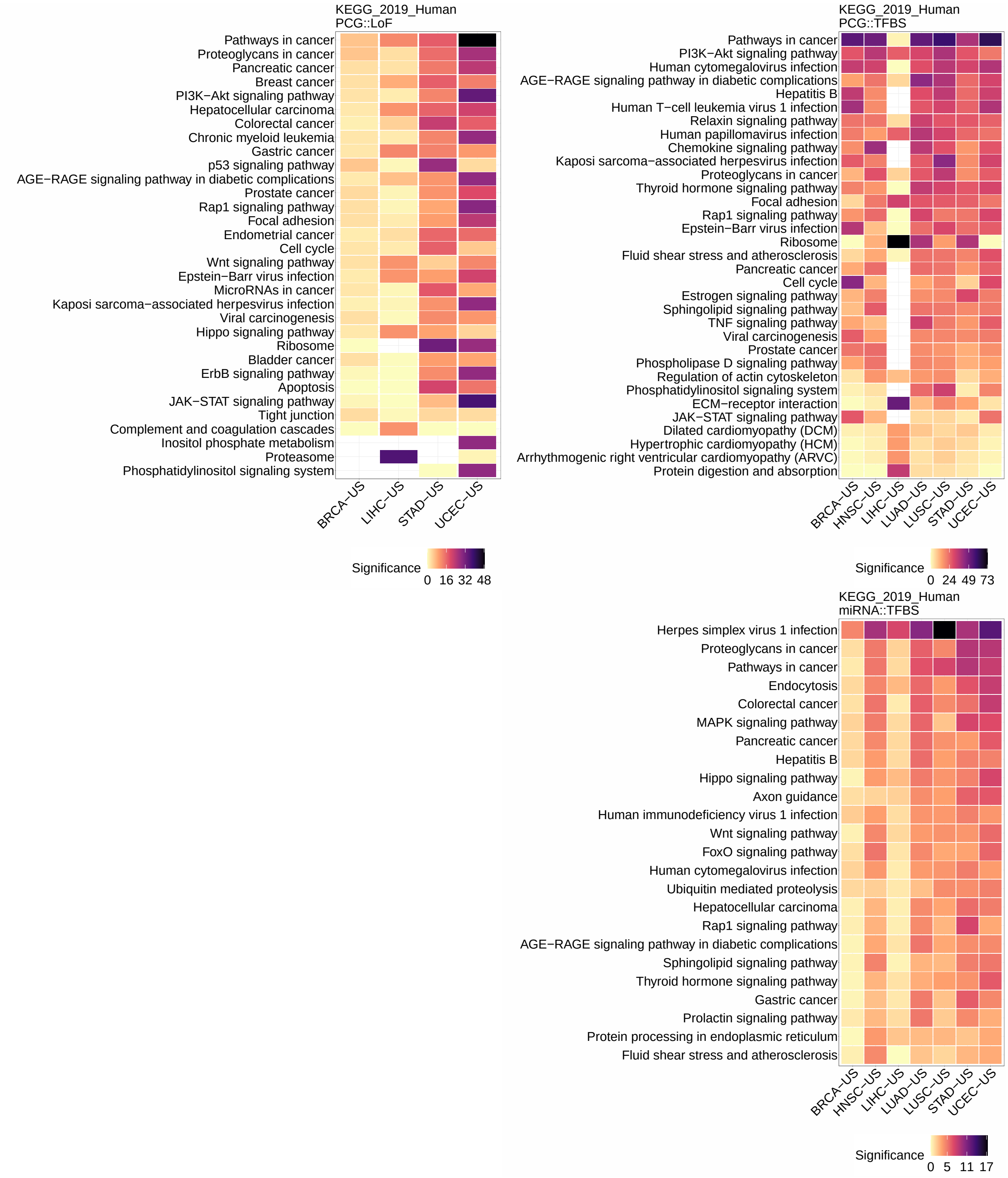


Figure S16: Functional enrichment (KEGG 2019) analysis considering dysregulated genes in the networks of predicted miRNA and protein-coding genes. The heatmaps represent the significance (-$\log_{10}$(P-value)) of the most enriched terms (rows) associated to the dysregulated genes found on the seven analyzed cohorts (columns) when considering protein-coding genes with LoF mutations (PCGs::LoF; top-left), protein-coding genes with cis-regulatory mutations (PCG::TFBS; top-right), and miRNAs with cis-regulatory mutations (miRNAs::TFBS; bottom-right). The terms are ordered by their mean ranks across the cohorts.

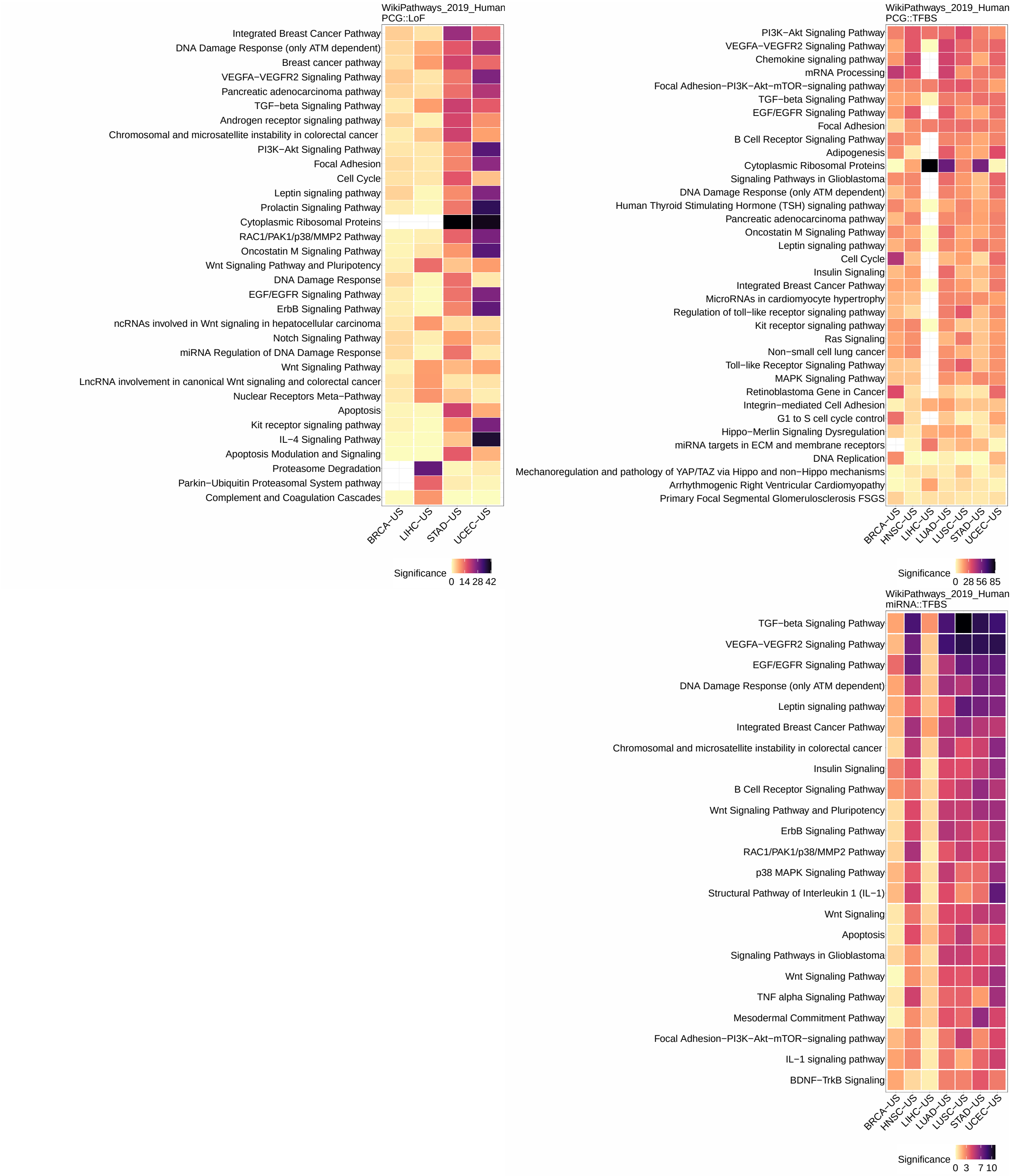


Figure S17: Functional enrichment (WikiPathways 2019 human) analysis considering dysregulated genes in the networks of predicted miRNA and protein-coding genes. The heatmaps represent the significance (-$\log_{10}$(P-value)) of the most enriched terms (rows) associated to the dysregulated genes found on the seven analyzed cohorts (columns) when considering protein-coding genes with LoF mutations (PCGs::LoF; top-left), protein-coding genes with cis-regulatory mutations (PCG::TFBS; top-right), and miRNAs with cis-regulatory mutations (miRNAs::TFBS; bottom-right). The terms are ordered by their mean ranks across the cohorts.

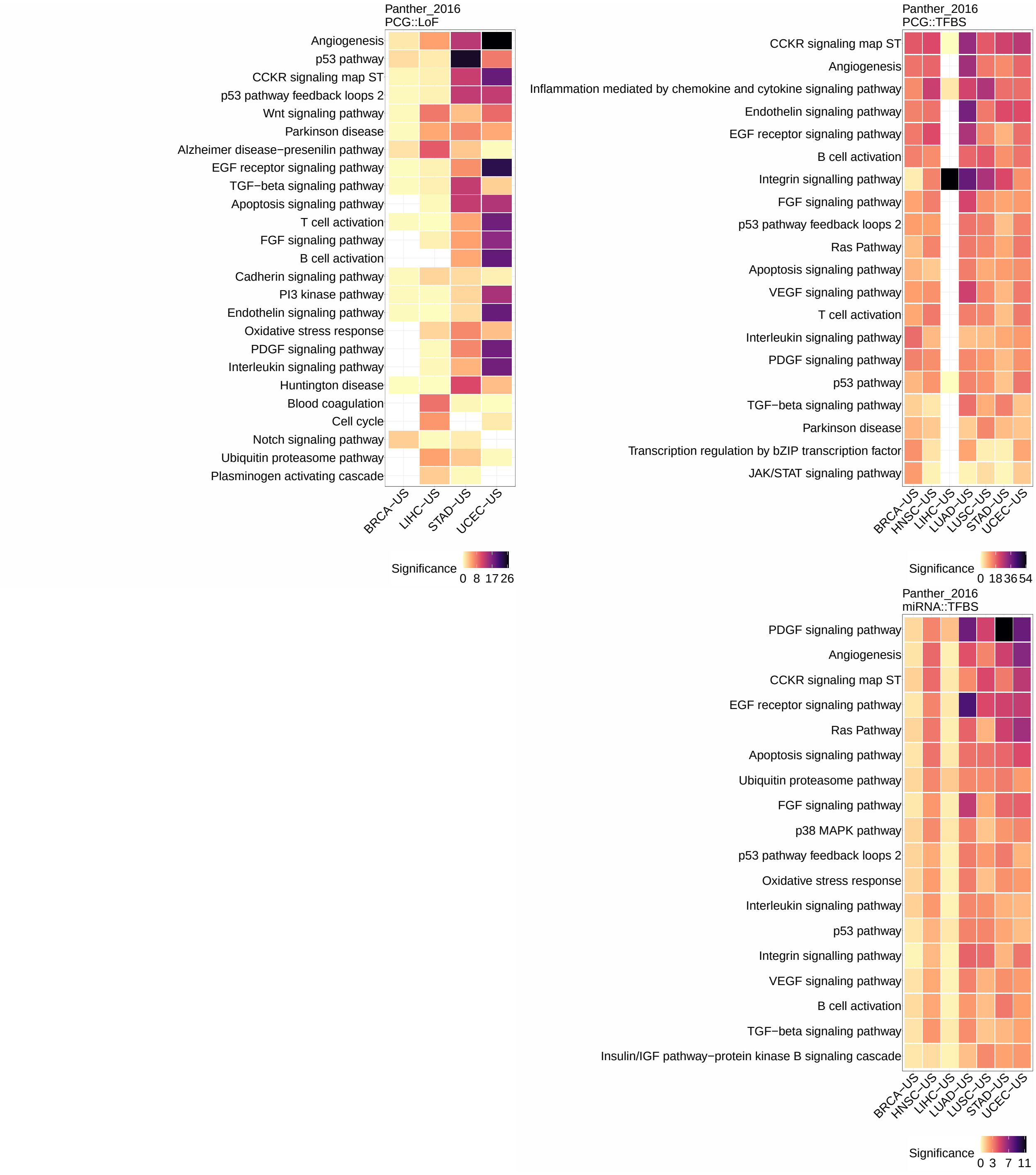


Figure S18: Functional enrichment (Panther 2016) analysis considering dysregulated genes in the networks of predicted miRNA and protein-coding genes. The heatmaps represent the significance (-log$_{10}$(P-value)) of the most enriched terms (rows) associated to the dysregulated genes found on the seven analyzed cohorts (columns) when considering protein-coding genes with LoF mutations (PCGs::LoF; top-left), protein-coding genes with cis-regulatory mutations (PCG::TFBS; top-right), and miRNAs with cis-regulatory mutations (miRNAs::TFBS; bottom-right). The terms are ordered by their mean ranks across the cohorts.

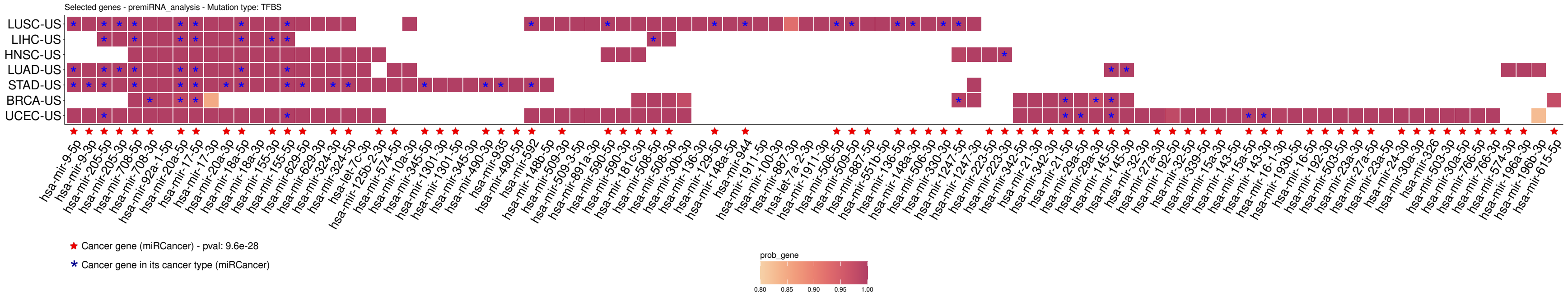


Figure S19: miRNAs predicted by xseq in the seven TCGA cohorts. Cell colors indicate the posterior probability computed over the corresponding cohort. Red stars indicate that the miRNA is annotated as a cancer miRNA in miRCancer. Blue stars indicate that the miRNA was reported as a cancer miRNA in the specific cancer type where it is predicted by xseq, according to miRCancer annotation.

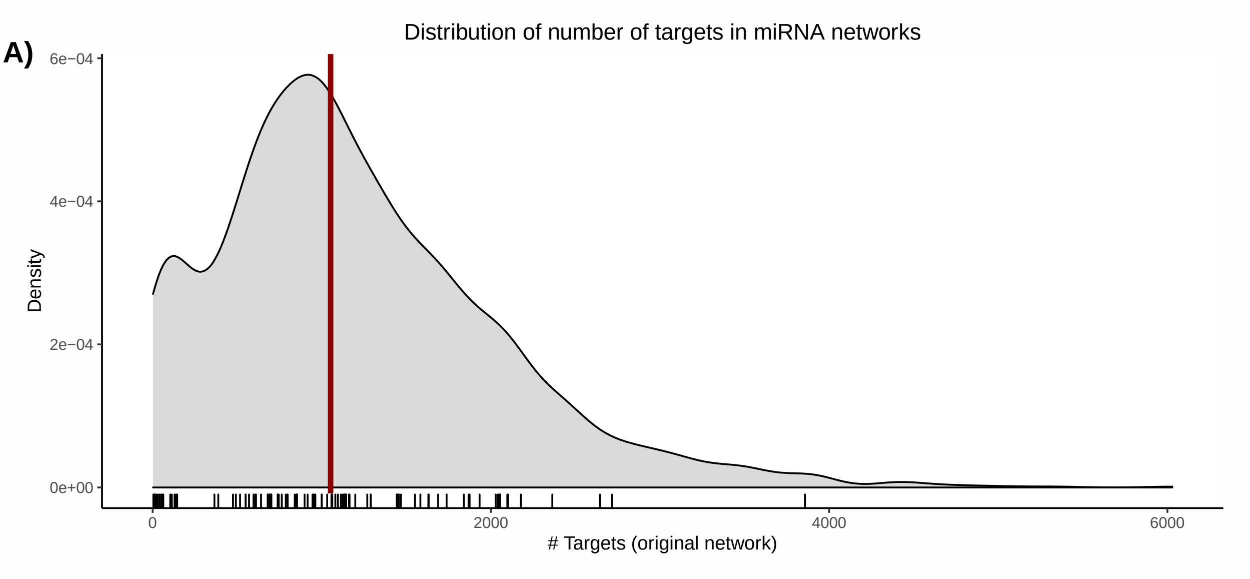


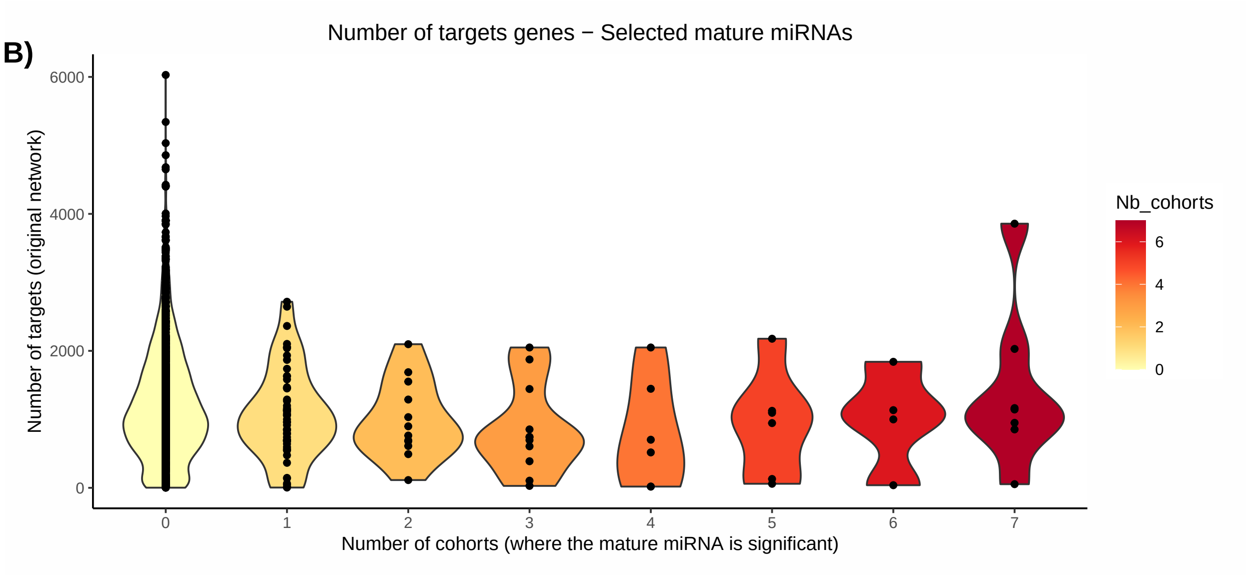


Figure S20: miRNA-target genes information. A) Distribution of the number of targets considered per miRNA. The red line indicates the median value (1052 targets). The rug at the bottom highlights the predicted miRNAs. B) Number of cohorts (x-axis) where each mature miRNA was predicted versus its number of targets (y-axis).

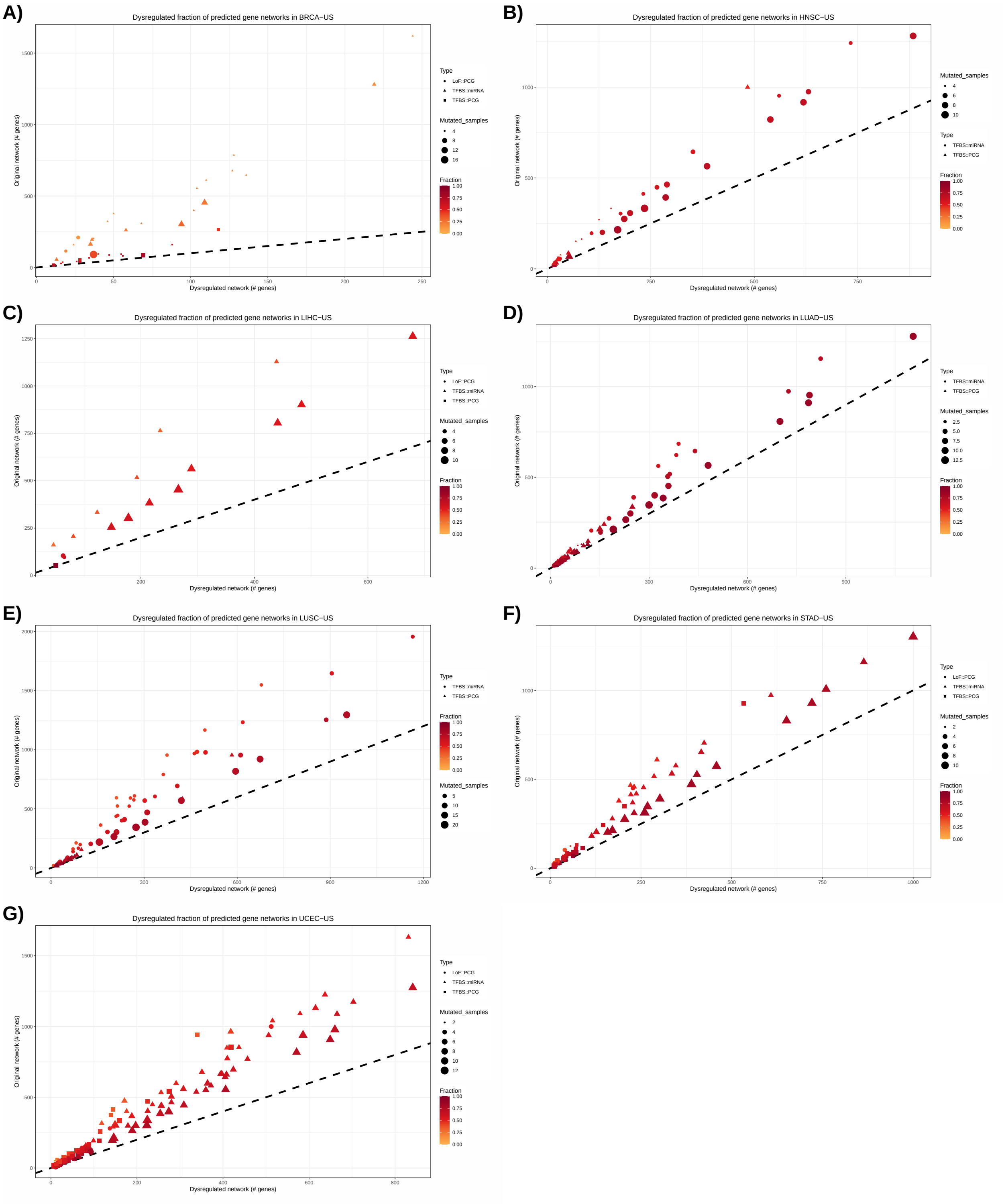


Figure S21: Fraction of the original gene networks predicted as dysregulated for the predicted genes in A) BRCA-US, B) HNSC-US, C) LIHC-US, D) LUAD-US, E) LUSC-US, F) STAT-US, and G) UCEC-US. X-axis corresponds to the number of dysregulated genes in the filtered network (see Material and methods), Y-axis corresponds to the original size of the gene networks before any filtering (for expression). Color scale corresponds to the fraction of the network that is predicted to be dysregulated in the samples with corresponding mutations of interest.

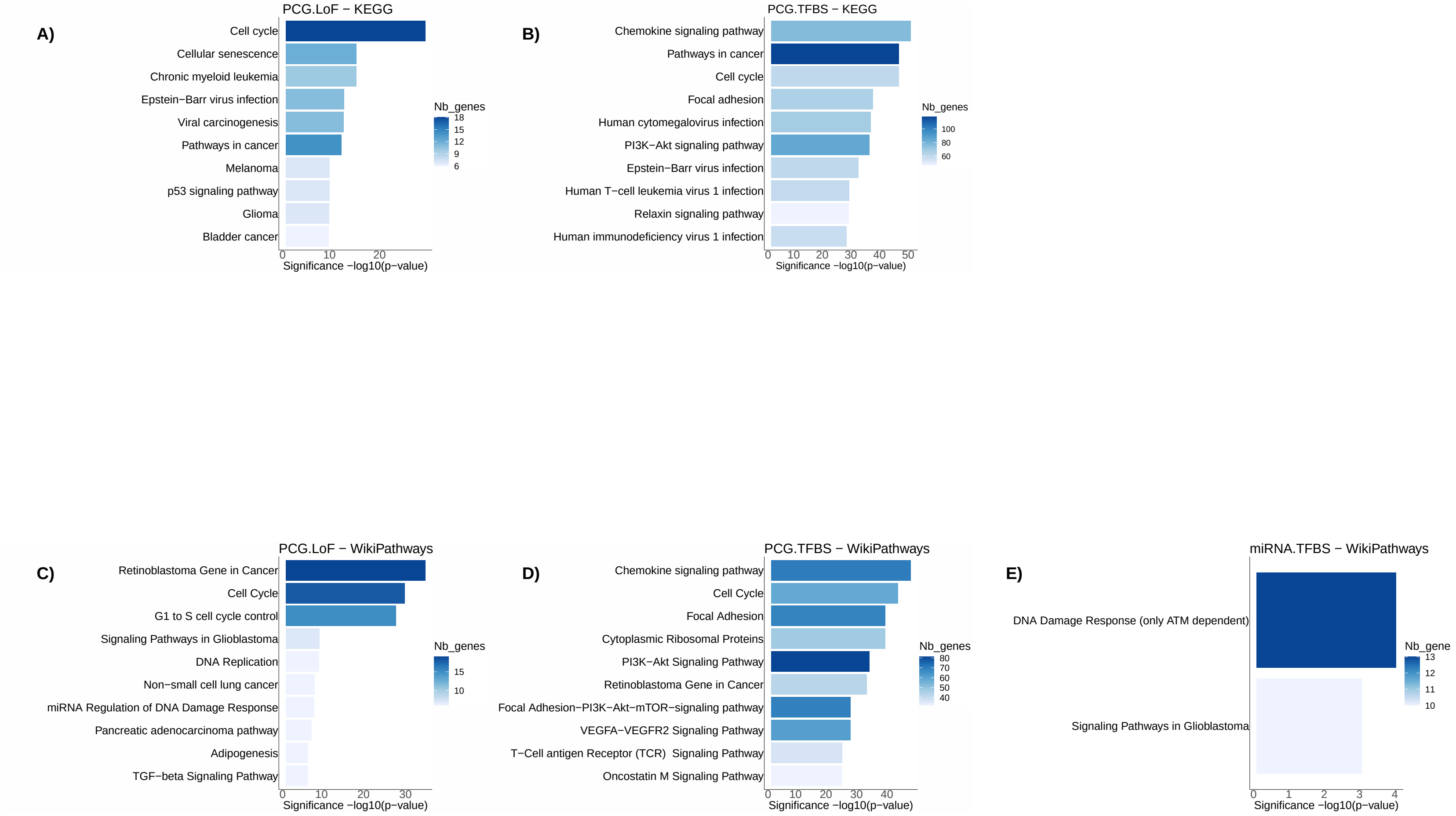


Figure S22: Functional enrichment analysis considering dysregulated genes in the networks of predicted miRNA and protein-coding genes in the BASIS cohort. The barplots represent the significance (-$\log_{10}$(P-value)) of the top-10 most enriched terms (rows) associated to the dysregulated genes when considering protein-coding genes with LoF mutations (PCGs::LoF; A-C), protein-coding genes with cis-regulatory mutations (PCG::TFBS; B-D), and miRNAs with cis-regulatory mutations (miRNAs::TFBS; E). Enriched terms from KEGG 2019 Human (A-B) and WikiPathways (C-E) are provided.

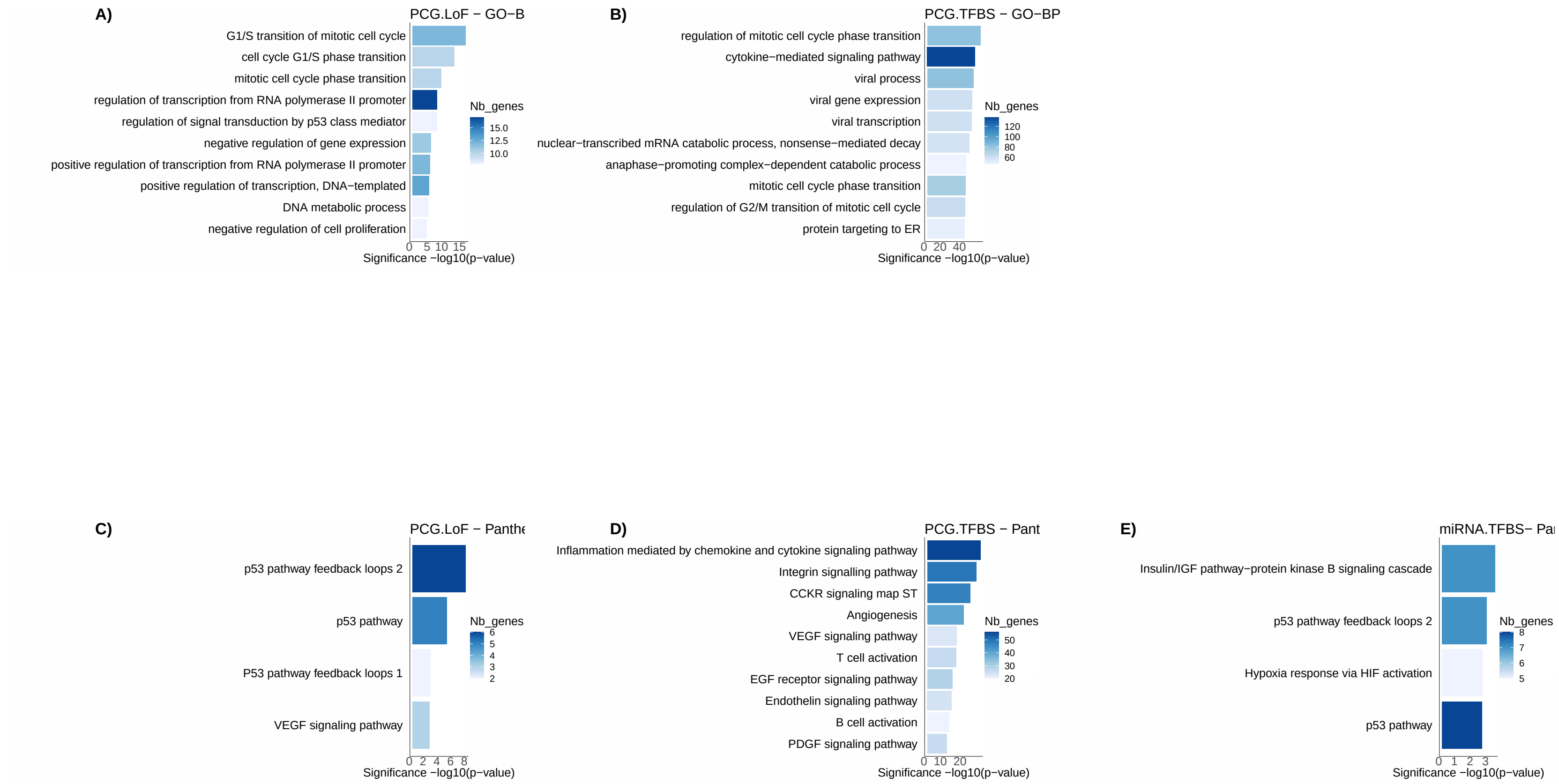


Figure S23: Functional enrichment analysis considering dysregulated genes in the networks of predicted miRNA and protein-coding genes in the BASIS cohort. The heatmaps represent the significance (-$\log_{10}$(P-value)) of the top-10 most enriched terms (rows) associated to the dysregulated genes when considering protein-coding genes with LoF mutations (PCGs::LoF; A-C), protein-coding genes with cis-regulatory mutations (PCG::TFBS; B-D), and miRNAs with cis-regulatory mutations (miRNAs::TFBS; E). Enriched terms from GO biological processes (GO-BP; A-B) and Panther 2016 (C-E) are provided.

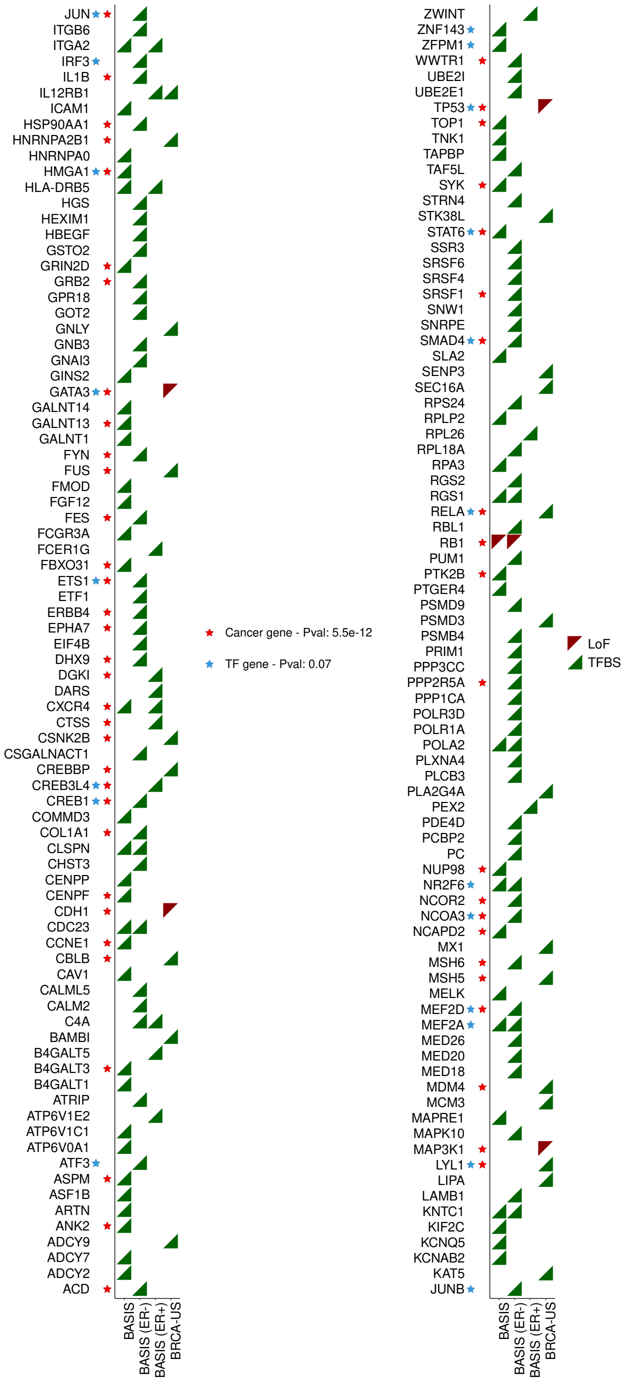


Figure S24: Predicted protein-coding genes in the BASIS cohort using all samples, ER+ samples and ER- samples, and in the TCGA BRCA-US cohort (columns). Predictions were obtained applying the xseq tool on each dataset independently considering protein-coding genes mutated through either LoF (red triangles) or cis-regulatory (TFBS; green triangles) mutations. Genes known as cancer genes (red stars) and TFs (blue stars) are highlighted and hypergeometric tests p-values for enrichment are provided in the legend (Material and methods).

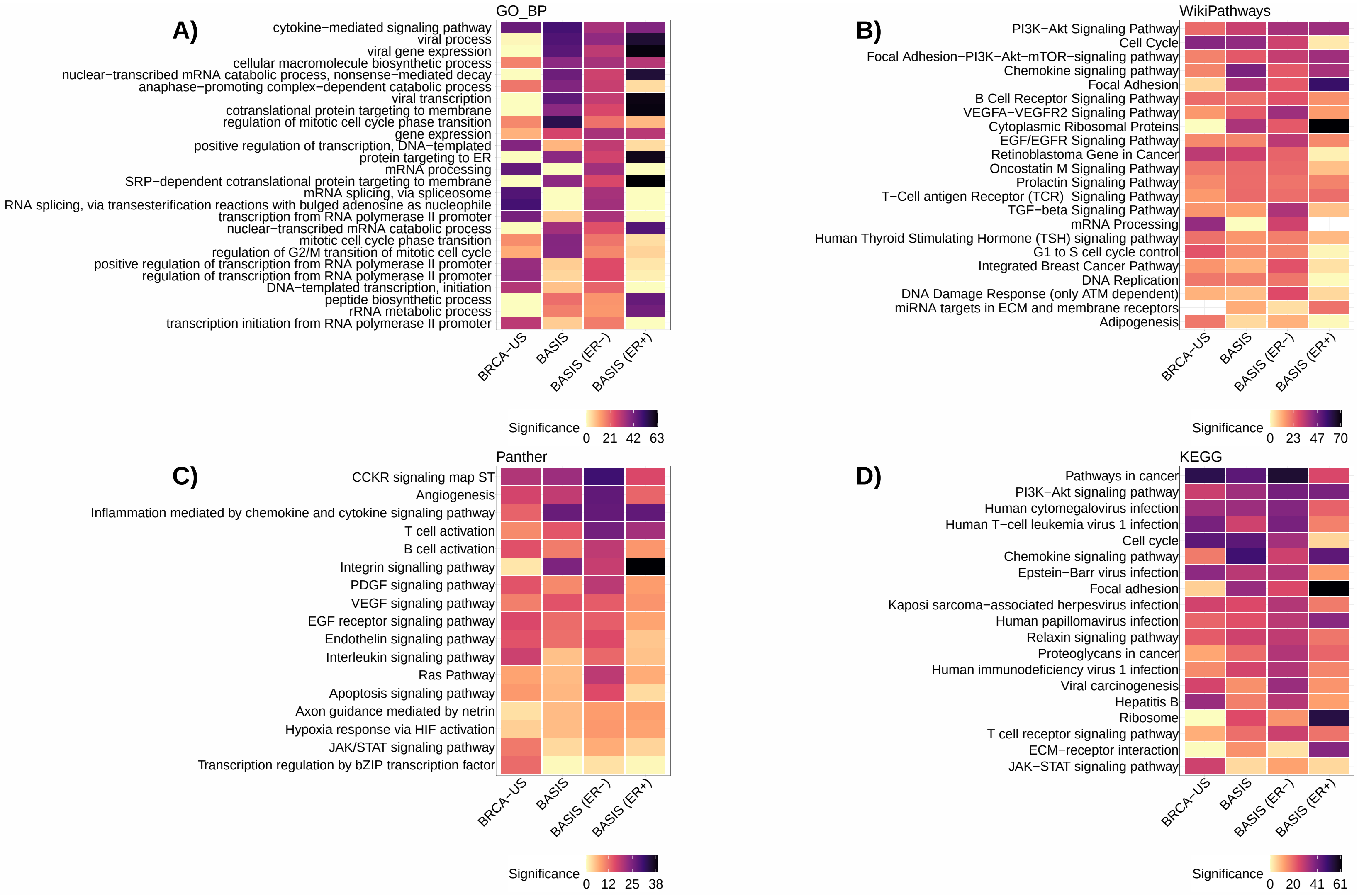


Figure S25: Functional enrichment analysis considering dysregulated genes in the networks of predicted protein-coding genes in the breast cancer cohorts (columns). The heatmaps represent the significance (-$\log_{10}$(P-value)) of the top-10 most enriched terms (rows) associated to the dysregulated genes for GO biological processes (GO-BP; A), WikiPathways (B), Panther (C), and KEGG (D). Terms (rows) are ordered by their mean rank across all cohorts.

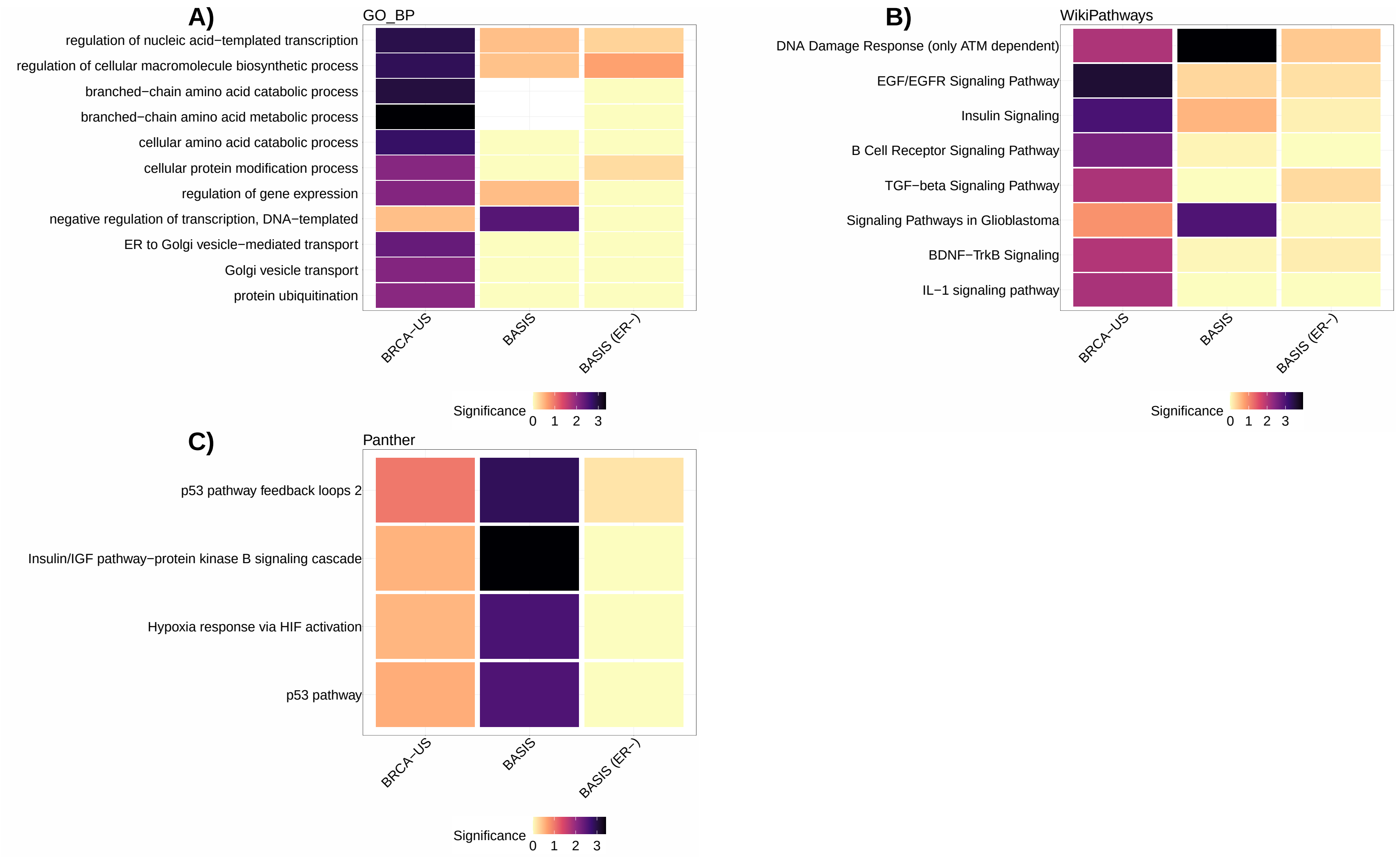


Figure S26: Functional enrichment analysis considering dysregulated target genes of predicted miRNAs in the breast cancer cohorts (columns). The heatmaps represent the significance (-$\log_{10}$(P-value)) of the top-10 most enriched terms (rows) associated to the dysregulated genes for GO biological processes (GO-BP; A), WikiPathways (B), and Panther (C). Terms (rows) are ordered by their mean rank across all cohorts.

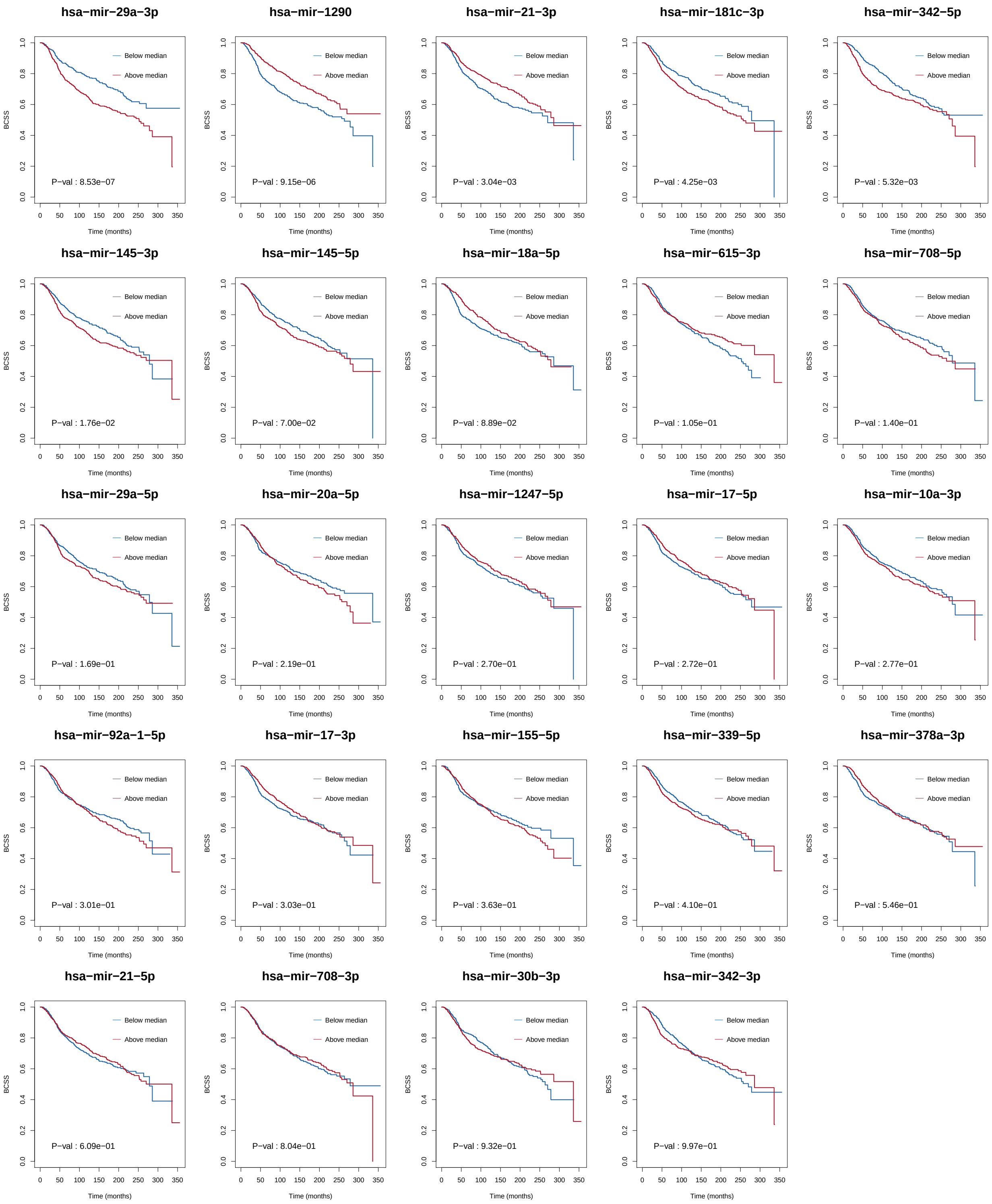


Figure S27: Breast cancer specific survival (BCSS, y-axis) plots for selected miRNAs. Kaplan-Meier survival curves were calculated using the METABRIC cohort. Samples were separated into two groups according to the level of miRNA expression (above/below the median).

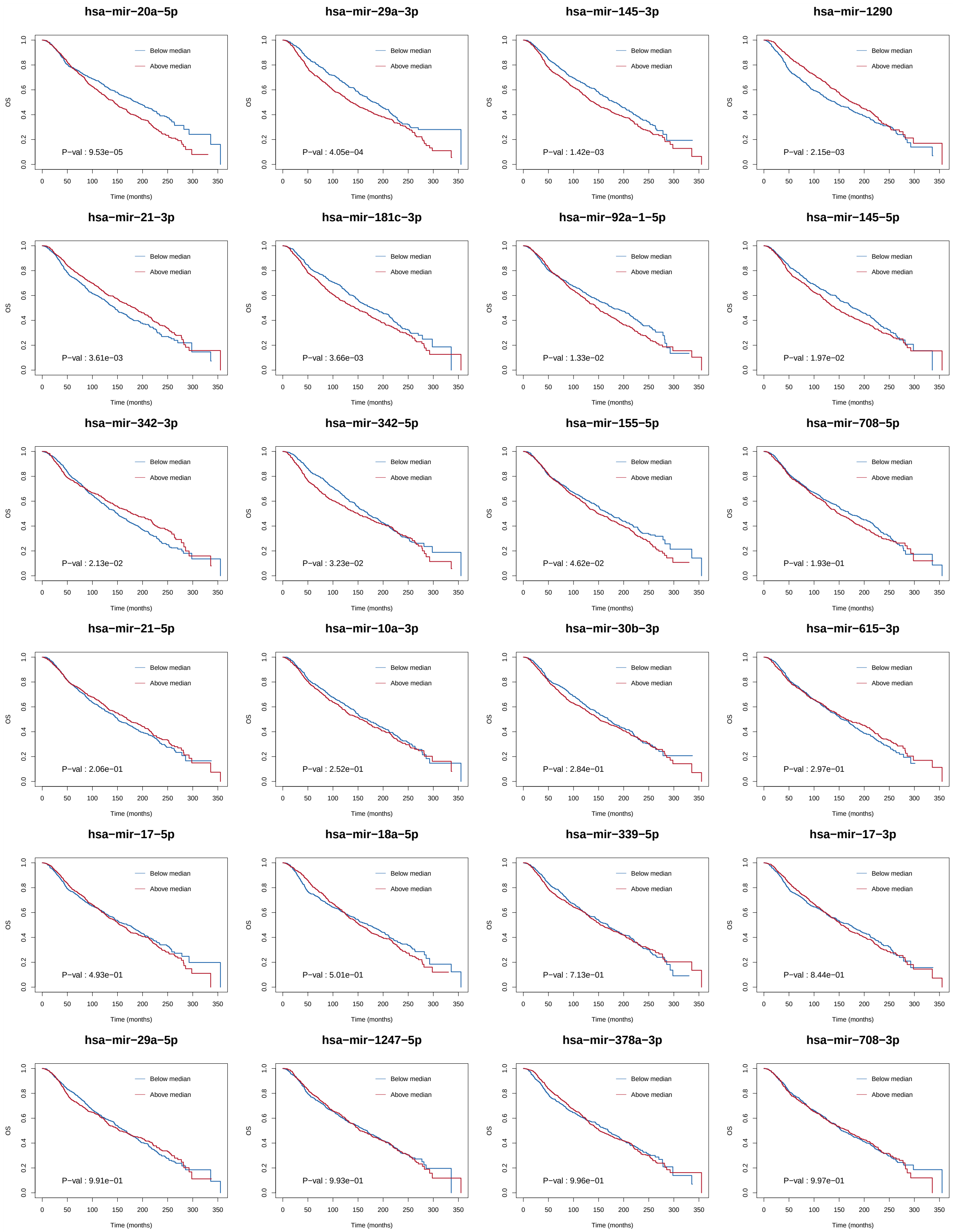


Figure S28: Overall survival (OS, y-axis) plots for selected miRNAs. Kaplan-Meier survival curves were calculated using the METABRIC cohort. Samples were separated into two groups according to the level of miRNA expression (above/below the median).
